# Basis of narrow-spectrum activity of fidaxomicin on gut pathogen *Clostridioides difficile*

**DOI:** 10.1101/2022.01.17.476619

**Authors:** Xinyun Cao, Hande Boyaci, James Chen, Yu Bao, Robert Landick, Elizabeth A. Campbell

## Abstract

Fidaxomicin (Fdx) is widely used to treat *Clostridioides difficile* (*Cdiff*) infections (CDIs), but the molecular basis of its narrow-spectrum activity in the human gut microbiome remains enigmatic. CDIs are a leading cause of nosocomial deaths. Fdx, which inhibits RNA polymerase (RNAP), targets *Cdiff* with minimal effects on gut commensals, reducing CDI recurrence. Here, we present the cryo-electron microscopy structure of *Cdiff* RNAP in complex with Fdx, allowing us to identify a crucial Fdx-binding determinant of *Cdiff* RNAP that is absent in most gut microbiota like Proteobacteria and Bacteroidetes. By combining structural, biochemical, and bioinformatic analyses, we establish that a single RNAP residue is a sensitizing element for Fdx narrow-spectrum activity. Our results provide a blueprint for targeted drug design against an important human pathogen.

## Main

*Clostridioides difficile* (*Cdiff*) is a gram-positive, spore-forming, and toxin-producing intestinal bacterium that infects the human gut and causes lethal diarrhea (*Cdiff* infections or CDIs). With the alarming increase in infections caused by highly pathogenic variants, *Cdiff* has been designated an “urgent threat” by CDC^1^. Broad-spectrum antibiotics like vancomycin and metronidazole are used to treat CDIs, but these antibiotics decimate the normal gut microbiome, paradoxically priming the gastrointestinal tract to become more prone to CDI recurrences^2,3^ (Fig.1a). In 2011, the macrocyclic antibiotic fidaxomicin (Fdx; Fig. 1b) became available to treat CDI. Fdx selectively targets *Cdiff* but does not effectively kill crucial gut commensals such as Bacteroidetes^4^, abundant microbes in the human gut microbiome that protects against *Cdiff* colonization^5,6^. Fdx targets the multisubunit bacterial RNA polymerase (RNAP, subunit composition α_2_ββ′ω), which transcribes DNA to RNA in a complex and highly regulated process. However, no structure is available for Clostridial RNAP. Studies using *Mycobacterium tuberculosis* (*Mtb*) and *Escherichia coli* (*Eco*) RNAPs show that Fdx functions by inhibiting initiation of transcription^7-10^. RNAP forms two mobile pincers that surround DNA^11,12^ and Fdx inhibits initiation by jamming the pincers in an “open” state, preventing one pincer, the clamp, from closing on the DNA. This doorstop-like jamming results in failures both to melt promoter DNA and to secure the DNA in the enzyme’s active-site cleft. Although the general architecture of RNAP is similar for all cellular organisms, differences in the primary subunit sequences, peripheral subunits, or lineage-specific insertions that occur in bacterial RNAP^13^ could explain Fdx sensitivity. For example, *Mtb* RNAP is much more sensitive to Fdx than *Eco* RNAP^7^, and the essential transcription factor RbpA sensitizes *Mtb* to Fdx even further^7^. The half-maximal inhibitory concentration or IC50 is 0.2 μM for *Mtb* RNAP with full-length RbpA, 7 μM for *Mtb* RNAP lacking the RbpA–Fdx contacts, and 53 μM for *Eco* RNAP (Extended Data Table 1)^7^. However, *Cdiff* lacks RbpA leaving the molecular basis of *Cdiff* sensitivity unresolved.

**Fig. 1.**
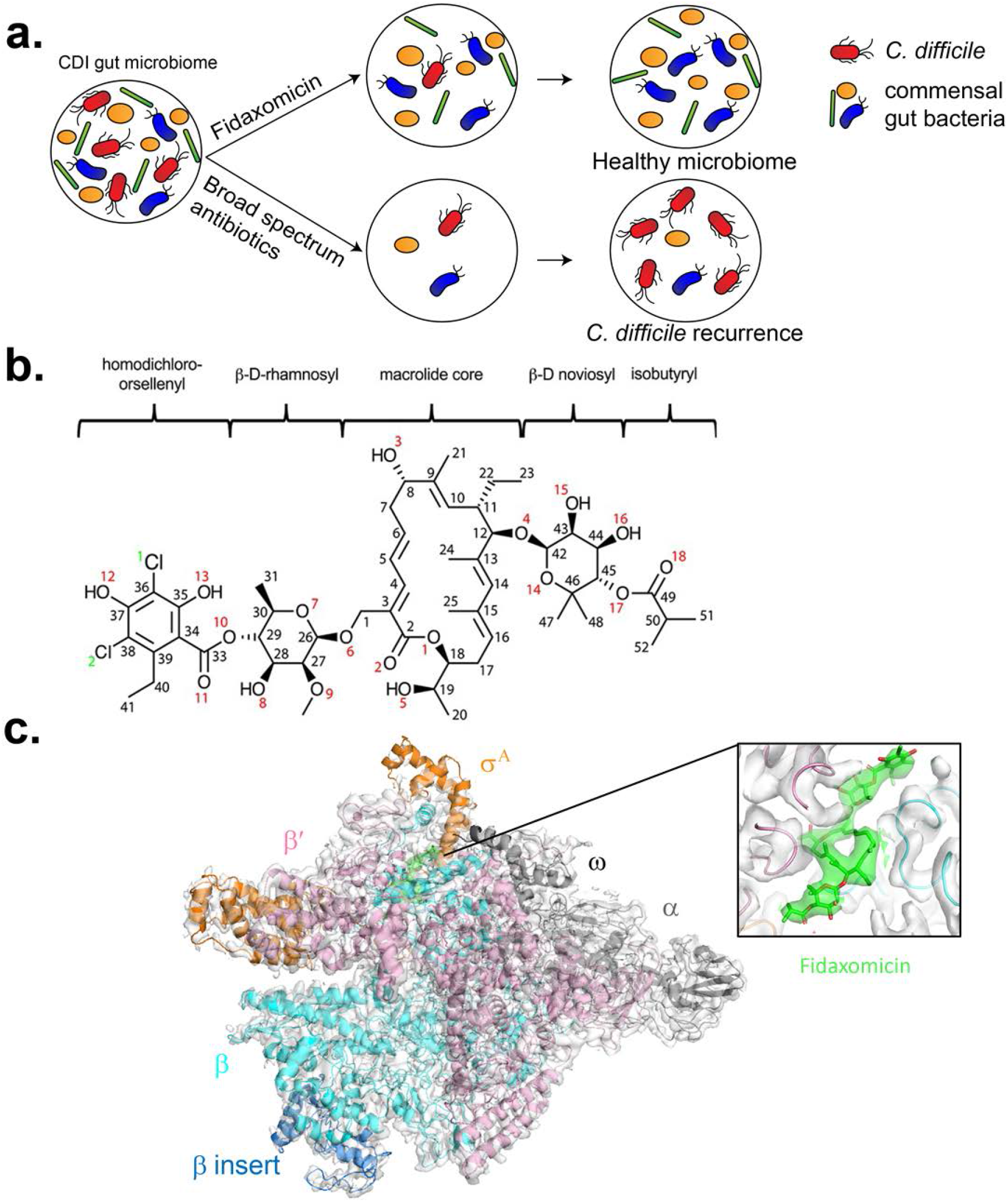
Fidaxomicin is a narrow-spectrum antimicrobial that inhibits RNAP. **a**, Diagram illustrating how fidaxomicin specifically targets *Cdiff* without affecting gut commensals and thus reduces recurrence (upper circles). For CDI patients treated with broad-spectrum antibiotics (lower circles), the abundance of gut commensals drops simultaneously with *Cdiff* resulting in high rates of *Cdiff* recurrence. **b**, Chemical structure of Fdx. **c**, Cryo-EM structure of *Cdiff* Eσ^A^ in complex with Fdx. The Eό^A^ model is colored by subunits according to the key, and the 3.26 Å cryo-EM map is represented in a white transparent surface. The cryo-EM density for Fdx is shown in the inset as a green transparent surface.

The pathogenicity and limited genetic tools for *Cdiff* complicate using *Cdiff* directly for structural and mechanistic studies of RNAP, with a single report of endogenous *Cdiff* RNAP purification yielding small amounts of enzyme with suboptimal activity^14^. To enable study of *Cdiff* RNAP, we created a recombinant system in *Eco* that yields milligram quantities of *Cdiff* core RNAP (E) and also enables rapid mutagenesis (see methods and supplement). *Cdiff* housekeeping σ^A^ factor was also expressed in *Eco*, purified, and combined with core *Cdiff* RNAP to produce the holoenzyme (Eσ^A^). The purity, activity, and yield of Eσ^A^ were suitable for structural and biochemical studies (Extended Data Fig. 1a, 1b and 1c).

To visualize the binding of Fdx to its clinical target, we used single particle cryo-electron microscopy (cryo-EM) to solve the structure of *Cdiff* Eσ^A^ in complex with Fdx. We obtained a cryo-EM map representing a single structural class comprising *Cdiff* Eσ^A^ and bound Fdx at 3.3 Å nominal resolution, with a local resolution of ∼2.7-3 Å around the Fdx-binding pocket (Fig. 1c, Extended Data Figs. 2 and 3, Extended Data Table 2)^15^. The structure reveals key features of the *Cdiff* Eσ^A^ and provides the first view of a Clostridial Eσ^A^.

*Cdiff* RNAP contains a lineage-specific insert in the β lobe domain that resembles one found in RNAP from *Bacillus subtilis* (*Bsub*, a Firmicute like *Cdiff*), but distinct from the better-characterized inserts in *Eco* RNAP^16-18^. The Firmicute β insert corresponds to βi5 identified by sequence analysis^13^ and consists of two copies of the β-β’ module 2 (BBM2) protein fold whereas the β lobe insert in *Eco* RNAP occurs at a different position and corresponds to βi4 (Extended Data Fig. 4). Our structure revealed *Cdiff* βi5 (4-5 Å; Extended Data Fig. 3) at a position similar to *Bsub* βi5 ^17^ but the *Cdiff* insert is larger (121 amino acids vs 99 amino acids in *Bsub*) (Extended Data Fig. 4). The function of the Firmicute βi5 is unknown and awaits further study but is unlikely to impact Fdx binding (located ∼70 Å away) or activity.

*Cdiff* σ^A^ possesses all conserved regions of σ, which were located in the *Cdiff* Eσ^A^ structure at locations similar to those seen for other bacterial housekeeping σ-factors (Extended Data Fig.5)^7,19,20^. Cryo-EM density was not visible for most of σ region 1 (residues 1–115 of the 150 residues in *Cdiff* region 1), as also seen with other bacterial holoenzymes characterized structurally^7,20,21^. *Cdiff* σ^A^ lacks the non-conserved region (NCR) insert between regions and 1 and 2 found in some other bacteria like *Eco* (Extended Data Fig. 5)^16^.

As seen in other RNAPs^7,8,22^, Fdx appears to stabilize the clamp pincer in an open state, but the *Cdiff* clamp is twisted slightly toward Fdx relative to the *Mtb* Eσ^A^ structure with Fdx (Fig.2a). Opening and closing of RNAP’s pincers are required for transcription initiation^11,22^. Fdx binds *Mtb* RNAP at a hinge between two RNAP pincers (the β′ clamp and β lobe pincers), thus physically jamming the hinge and locking RNAP in an open conformation that is unable to form a stable initiation complex^8,9^. Fdx occupies the same location in the *Cdiff* Eσ^A^–Fdx structure, indicating that the hinge-jamming mechanism is widely conserved. However, the slight twisting of the *Cdiff* β′ clamp pincer relative to that observed in Fdx-bound *Mtb* RNAP (Fig. 2a) increases clamp–Fdx contacts (see below).

**Fig. 2.**
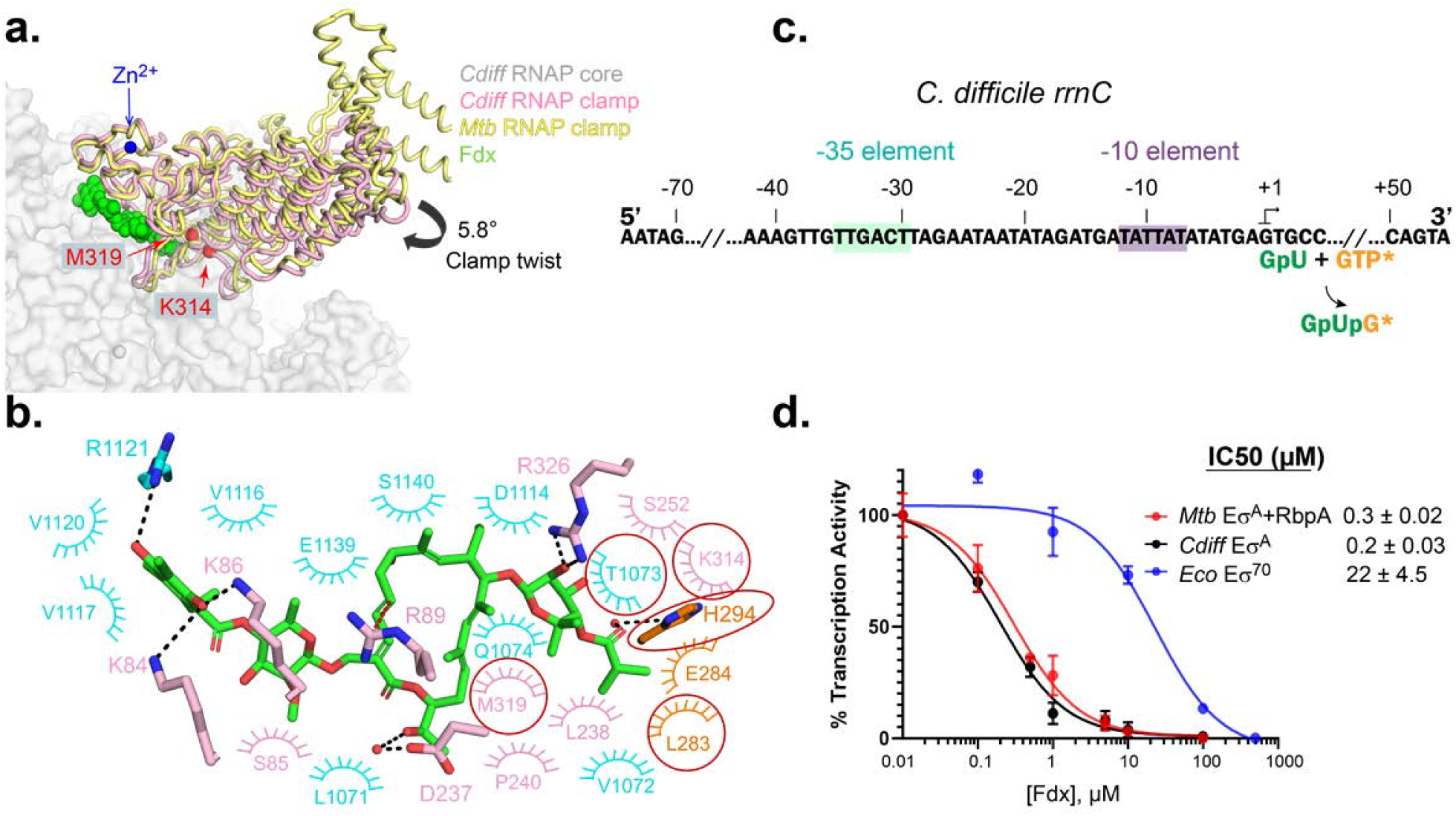
Fdx binding and inhibition of the *Cdiff* Eσ^A^. **a**, Clamp differences between *Cdiff* and *Mtb* RNAP. The *Cdiff* RNAP clamp (pink) is twisted (5.8°) towards Fdx compared with the *Mtb* RNAP–Fdx clamp (yellow) (PDB ID: 6BZO). The actinobacterial specific insert in the clamp is partially cropped. The clamp residues, β′K314 and β′M319, that interact with Fdx (shown in green spheres) in *Cdiff* but not *Mtb* RNAP are shown as red spheres. The zinc in the ZBD is shown as a blue sphere. **b**, The interactions between Fdx and *Cdiff* RNAP are shown. Hydrogen-bonding interactions are shown as black dashed lines. The cation-π interaction of β′R89 is shown with a red dashed line. Arches represent hydrophobic interactions. RNAP residues are colored corresponding to subunits: cyan (β) and pink (β′). The Fdx-contacting residues that are not present in the *Mtb* Eσ^A^–Fdx structure (pdb 6BZO) are marked with red circles. **c**, The sequence of the native *Cdiff* ribosomal RNAP *rrnC* promoter used in the *in vitro* transcription assay in **d**. The –10 and –35 promoter elements are shaded in purple and green, respectively. The abortive transcription reaction used to test Fdx effects is indicated below the sequence (*, [α-^32^P]GTP used to label the abortive product GpUpG). **d**, Fdx inhibits *Cdiff* Eσ^A^ and *Mtb* RbpA-Eσ^A^ similarly and ∼100 times more effectively than *Eco* Eσ^70^. Error bars are standard deviations (SD) of three independent replicates (for some points, SD was smaller than the data symbols).

Fdx contacts six key structural components of *Cdiff* RNAP: β clamp, β′ switch region 2 (SW2), β switch region 3 (SW3), β switch region 4 (SW4), β′ zinc-binding-domain (ZBD), and β′ lid (Figs. 2b and 3). We compared the Fdx-binding determinants in *Cdiff* RNAP to those previously determined in *Mtb* RNAP^7,8^. Most of the interactions between Fdx and RNAP were conserved between the two species (Extended Data Fig. 6, Extended Data Table 3). In *Cdiff* RNAP (*Mtb* numbering in parentheses), Fdx formed direct hydrogen bonds or salt bridges with four residues βR1121 (K1101), β′K84 (R84), β′K86 (K86), and β′R326 (R412) and two water-mediated hydrogen bonds with β′D237 (E323) and σH294 (Q434). (Figs. 2b, Extended Data Fig. 6, and Extended Data Table 3). Fdx binding is also stabilized by a cation-π interaction between the β′R89 and the Fdx macrolide core C3–C5 double bond in both *Mtb* and *Cdiff* RNAPs. *Cdiff* RNAP residues known to confer Fdx-resistance when mutated (βV1143G, βV1143D, βV1143F, β′D237Y and βQ1074K^23,24^) were located within 5 Å of Fdx (Extended Data Fig. 6).

Of particular interest, β′K84 in *Cdiff* RNAP forms a salt bridge with the oxygen on the phenolic group of Fdx (due to the acidity of phenol) whereas the corresponding residue (β′R84) in *Mtb* forms a cation-π interaction with the aromatic ring of the Fdx homodichloroorsellinic acid moiety (Fig. 2b). We propose that these coulombic interactions by β′K84 (β′R84) sensitize both RNAPs to tight Fdx binding (see comparison of individual residues in *Cdiff* and *Mtb* RNAPs that bind Fdx; Extended Data Figs. 6 and 7, Extended Data Table 3). *Mtb* RbpA, an essential transcriptional regulator in mycobacteria, lowers the IC50 of Fdx by a factor of 35 via Fdx contacts with two RbpA residues in the N-terminal region^7^. *Cdiff* RNAP lacks a RbpA homolog, but we observed four hydrophobic interactions between Fdx and *Cdiff* RNAP (with βT1073, β′M319, β′K314, and σL283) and one water-mediated hydrogen bonding interaction with σH294 that are not present with the corresponding *Mtb* RNAP residues (Fig. 2b, Extended Data Fig.6, Extended Data Table 3). Some of these new interactions (β′M319 and β′K314 in the clamp) with Fdx are created by the relatively increased rotation of the *Cdiff* RNAP clamp towards Fdx (Fig. 2a).

Gram-negative bacteria are more resistant to Fdx than gram-positive bacteria^25-27^. This dichotomy could reflect differences in membrane and cell-wall morphology, differences in RNAPs, or both. To compare the activity of Fdx against *Cdiff* and *Mtb* RNAP, we performed abortive transcription assays using purified RNAPs and the native *Cdiff rrnC* ribosomal RNA promoter (Fig. 2c)^14^. The IC50s of Fdx for *Cdiff* RNAP Eσ^A^ (∼0.2 μM) and *Mtb* Eσ^A^ including RbpA (∼0.3 μM) are similar, consistent with our structural observations (Fig. 2d, Extended Data Fig. 6). These IC50s are two orders of magnitude lower than that for *E. coli* σ^70^-holoenzyme (Eσ^70^) on the same DNA template (Fig. 2d, Extended Data Fig. 8), suggesting that the differences in RNAPs contribute significantly to the differences in MICs between *Cdiff* (a gram-positive) and gram-negative bacteria. This observation suggests that the Fdx-binding residues identified in *Mtb* and *Cdiff* RNAPs can be used as a reference to predict Fdx potency in other bacterial species, including gut commensals^25^.

We next used our *Mtb* and *Cdiff* RNAP–Fdx structures to predict the interactions responsible for the narrow spectrum activity of Fdx. Using sequence alignments of β′ and β from bacterial species with reported Fdx MICs^25,27^ (Extended Data Fig. 9), we found that the Fdx-binding residues identified in *Mtb* and *Cdiff* RNAP are mostly conserved among these divergent bacteria, except for the aforementioned β′K84 (β′R84 in *Mtb*) and β′S85 (β′A85 in *Mtb*). β′K84 and S85 are located in the ZBD. β′K84 forms a salt bridge with the likely ionized Fdx O13 whereas S85 Cβ forms a nonpolar interaction with the Fdx C32 methyl group (Figs. 1b, 3 and Extended Data Fig. 6, Extended Data Table 3). We focused on β′K84 because all species contain a Cβ at position 85 whereas position 84 displays an intriguingly divergent pattern among gut commensal bacteria (Extended Data Fig. 9). For gram-positive bacteria, which are hypersensitive to Fdx (MIC<0.125 εg/mL), the β′K84 position is always positively charged (K or R). However, for gram-negative bacteria, which are resistant to Fdx (MIC >32 εg/mL), β′K84 is replaced by a neutral residue (Q in *Eco* or L in *Pseudomonas aeruginosa* and *Neisseria meningitidis*; Extended Data Fig. 9). Notably, in Bacteroidetes, which are highly resistant to Fdx (MIC >32 εg/mL, e.g., *Bacteroides uniformis, Bacteroides ovatus*, and *Bacteroides distasonis*), β′K84 is replaced by negatively charged glutamic acid (E). In an analysis of common species present in the human gut microbiota (Extended Data Table 4)^28,29^, β′K84 is replaced by E in Bacteroidetes (the most abundant bacteria^30,31^) and by neutral residues (Q, T, or S) in Proteobacteria (Figs. 4a, 4b, and Extended Data Fig.10). We thus refer to β′K84 as the Fdx sensitizer and propose it is crucial for tight Fdx binding in two ways: first, by forming a salt bridge (a proton-mediated ionic interaction) between the positively charged ε-amino group of β′K84 and a negatively charged phenolic oxygen of Fdx; and second, by rigidifying the α-helix of the ZBD and thus facilitating backbone hydrophobic and hydrogen-bonding interactions with downstream residues S85 (A85 in *Mtb*) and K86 (Fig. 3 and Extended Data Figs. 6 and 7).

**Fig. 3.**
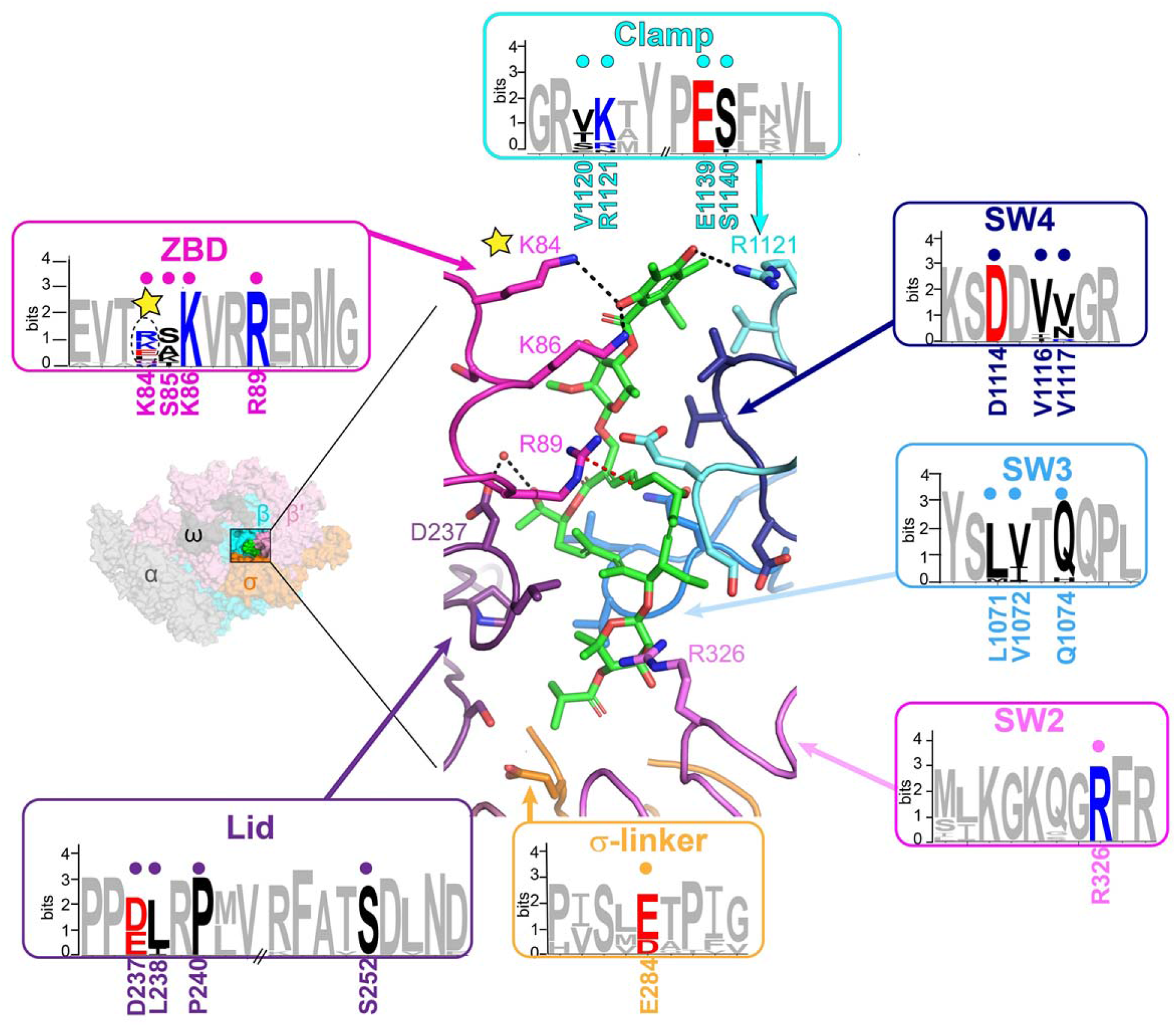
Analysis of Fdx-interacting residues across bacterial lineages. *Cdiff* RNAP Eσ^A^–Fdx is shown as a molecular surface for orientation (left inset). The boxed region is magnified on the right with RNAP subunits shown as α-carbon backbone worms, Fdx-interacting residues conserved between *Mtb* and *Cdiff* shown as side-chain sticks, and Fdx (green) shown as sticks. Non-carbon atoms are red (oxygen) and blue (nitrogen). The *Mtb*–*Cdiff*-conserved, Fdx-interacting residues shown in the cartoon structure are labeled under the sequence logos. Amino acids that make hydrophilic interactions with Fdx are labeled on the structure. Representative bacterial species with published Fdx MICs were used to make the logos^25^. See Extended Data Fig. 9 for detailed sequence alignments. Most Fdx-interacting residues are conserved except residues corresponding to *Cdiff* β′S85 and the sensitizer (yellow star corresponding to *Cdiff* β’K84).

**Fig. 4.**
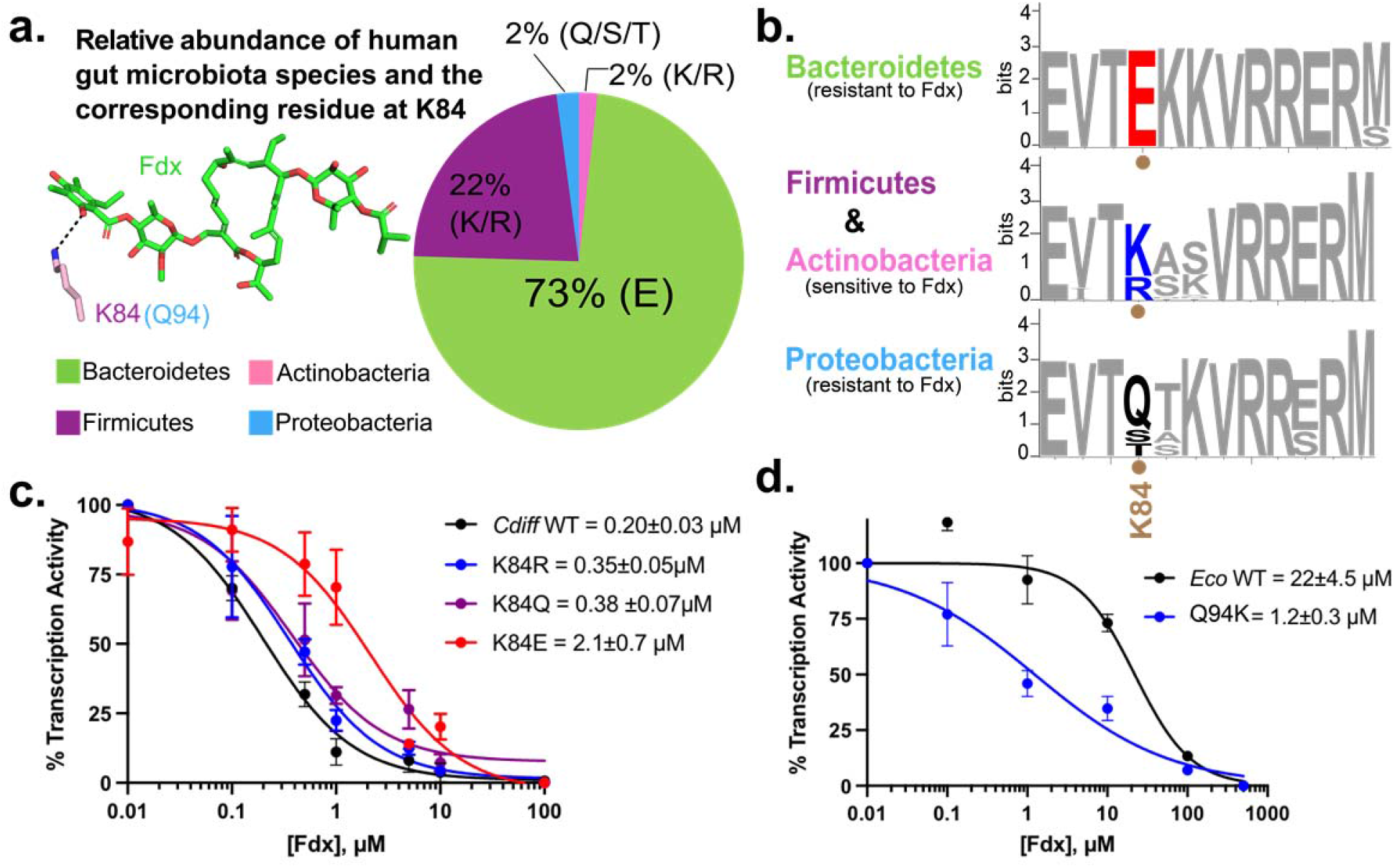
The sensitizer position (β′K84 in *Cdiff* RNAP) explains Fdx narrow-spectrum activity in the gut microbiota. **a**, The amino acid present at the sensitizer position among the four most common bacterial phyla in the human microbiota (specified after %) compared to the relative abundance of bacteria in each phylum shown in the pie chart as a percent of total microbiota (adapted from Refs.^31^ and^30^). The amino acids present at the sensitizer position for each phylum were identified based on 66 representative species^28 29^ (Extended Data Fig.10, Extended Data Table 4). **b**, Sequence logos for the Fdx-interaction region of the β′ZBD is highly conserved among Bacteroidetes, Firmicutes, Actinobacteria, and Proteobacteria except for the sensitizer position and C-adjacent residue (β′K84 and S85 in *Cdiff* RNAP). The logos were derived using the same 66 representative species (Extended Data Table 4). **c**, Fdx effects on abortive transcription reveal that the β’K84E substitution increases resistance 10-fold, whereas β′K84Q and β′K84R have much lesser effects. **d**, Transcription assays with *Eco* Eσ^70^ show the β′Q94K substitution in the *Eco* RNAP reduces Fdx IC50 by a factor of ∼20 relative to the WT enzyme.

We hypothesized that variation in the Fdx sensitizer plays a key role in determining the potency of Fdx activity on RNAP from different clades. To test this hypothesis, we constructed substitutions β′K84E, β′K84Q, and β′K84R in *Cdiff* RNAP and β′Q94K in *Eco* RNAP and then compared their inhibition by Fdx using the *Cdiff rrnC* abortive initiation assay (Figs. 4c, 4d, Extended Data Figs. 11, and 12). Fdx inhibits *Cdiff* wild-type (WT), β′K84Q, and β′K84R RNAPs at similar sub-μM concentrations. However, inhibition of *Cdiff* β′K84E RNAP requires a ten-fold higher concentration of Fdx than WT, indicating greater resistance to Fdx (Fig. 4c and Extended Data Fig.11). This result is consistent with our hypothesis that the negatively charged carboxyl group on the side chain of β′K84E repels to the negative oxygen of Fdx and disrupts the polar interaction, whereas the β′K84Q must be less disruptive to *Cdiff* RNAP–Fdx interaction possibly because the glutamine is capable of forming a hydrogen bond with Fdx.

To test the effect of positive versus neutral charge at the Fdx sensitizer in the context of an RNAP that is relatively Fdx-resistant, we compared WT and β′Q94K *Eco* RNAPs. The *Eco* β′Q94K substitution endows a positive charge at the position and dramatically increases sensitivity to Fdx (the IC50 decreased by a factor of 20; Fig. 4d, Extended Data Fig. 12). This result indicates that the lack of positive charge at the sensitizer position is indeed a crucial contributor to resistance to Fdx in Proteobacteria and posed an interesting discrepancy with the lack of effect of the *Cdiff* β′K84Q substitution. We hypothesize that other differences between *Cdiff* and *Eco* RNAP such as the relative flexibilities of the ZBD and clamp allow *Cdiff* RNAP, but not *Eco* RNAP, to sustain stronger interactions when the key position is neutral (Q). However, when a negative charge (E) is present at the sensitizer position, the repulsion between the carboxylic side chain (present in Bacteroidetes) and the Fdx phenolic oxygen leads to Fdx-resistance. The increased susceptibility of *Eco* β′Q94K RNAP to Fdx confirms that the positively-charged sensitizer (as found in *Cdiff* and *Mtb*) is crucial for conferring Fdx sensitivity.

In summary, our high-resolution structure of *Cdiff* RNAP Eσ^A^ reveals features that are likely to be specific to Clostridia and, to some extent Firmicutes and gram-positive bacteria. Analysis of this structure, in combination with bioinformatics and structure-guided functional assays, revealed a “sensitizing” determinant for Fdx, which turns out to be a single amino-acid residue in the ZBD of the RNAP β’ subunit. This work sheds light on how Fdx selectively targets *Cdiff* versus beneficial gut commensals like Bacteroidetes. Although wide-spectrum antibiotics are broadly effective therapies, our results highlight the advantages of narrow-spectrum antibiotics to treat intestinal infections and likely other bacterial infections. Treatment by narrow-spectrum antibiotics would reduce widespread antibiotic resistance and reduce the side effects caused by the collateral eradication of the beneficial bacteria in the gut microbiome. Using a similar approach to that applied here, further elucidation of diverse bacterial RNAP structures and mechanisms can provide a blueprint for designer antibiotics that leverage natural microbial competition to combat pathogens more effectively.

## Methods

Structural biology software was accessed through the SBGrid consortium^32^.

### Protein Expression and Purification

#### *Cdiff* σ^A^

The *Cdiff* σ^A^ gene (Kyoto Encyclopedia of Genes and Genomes (KEGG) entry: CD630_14550) was amplified from *Cdiff* 630 chromosomal DNA and cloned between NcoI and NheI sites of pET28a plasmid. A His_10_ tag with a Rhinovirus 3C protease recognition site was added to the N-terminus end of the σ^A^ gene to facilitate purification. *Escherichia coli (Eco)* BL21 (DE3) cells were transformed with this plasmid and were induced with 0.5 mM isopropyl-beta-D-thiogalactopyranoside (IPTG) overnight at 16 °C. The protein was affinity purified on a Ni^2+^-column (HiTrap IMAC HP, GE Healthcare Life Sciences). The eluted protein was cleaved with Rhinovirus 3C protease overnight and the cleaved complex was loaded onto a second Ni^2+^-column and the flow-through collected and further purified by size exclusion chromatography (Superdex 200, GE Healthcare) in buffer containing 20 mM Tris-HCl, pH 8, 5% (v/v) glycerol, 1 mM EDTA, 0.5 M NaCl, and 5 mM DTT. The eluted *Cdiff* σ^A^ were subsequently concentrated and stored at −80°C.

### *Cdiff* RNAP

The *Cdiff* RNAP overexpression plasmid was constructed in multiple steps. First, the *Cdiff 630 rpoA* (KEGG: CD630_00980), *rpoZ* (KEGG: CD630_25871), *rpoB* (KEGG: CD630_00660), and *rpoC* (KEGG: CD630_00670) genes were codon-optimized for *Eco* using Gene Designer (ATUM, Inc.) and codon frequencies reported by Welch et al^33^. A strong ribosome-binding site (RBS) was designed for each gene using the Salis RBS design tools (*https://www.denovodna.com/software/*)^34^. gBlock fragments containing *rpoA, rpoZ, rpoB and rpoC* genes and corresponding RBS were purchased from Integrated DNA Technologies directly. DNA fragments were assembled into pET21 (Novagen) using Gibson Assembly (NEB). Next, to maintain the subunit stoichiometry and prevent assembly with the host *Eco* subunits, β and β′ were fused using a polypeptide linker (LARHGGSGA). The same method was also used to construct overexpression plasmid for *Mtb* RNAP^35^. Finally, the His_10_ tag with a Rhinovirus 3C protease cleavable site was added to the C-terminus of *rpoC* to facilitate purification, resulting in plasmid pXC.026. The plasmids encoding *Cdiff* RNAP mutants (*rpoC* K84E, K84R, and K84Q) were constructed by Q5 site-directed mutagenesis (NEB) using pXC.026 as the template.

Overexpression of *Cdiff* RNAP yielded high levels of proteolysis and inclusion bodies in the conventional BL21 λDE3 strain. The yields of soluble *Cdiff* RNAP were increased in B834 (DE3), a strain reported to successfully produce gram-positive *Bacillus subtilis (Bsub)* RNAP^36^, The *Cdiff* core RNAP subunits were co-overexpressed in *Eco* B834 (Novagen) cells overnight at 16°C for ∼16 h after induction with 0.3 mM IPTG. The cell pellet was resuspended in the lysis buffer (50 mM Tris-HCl pH 8.0, 1 mM EDTA, 5% (v/v) glycerol, 5 mM 1,4-dithiothreitol (DTT), 1 mM protease inhibitor cocktail (PIC), and 1 mM phenylmethylsulfonyl fluoride (PMSF)). Cells were lysed by continuous flow through a French press (Avestin) and spun at 15,000 rpm twice for 20min each. Then the supernatant was then precipitated by the addition of 0.6% polyethyleneimine (PEI). PEI pellets were washed three times with a buffer containing 10 mM Tris-HCl, pH 8, 0.25 M NaCl, 0.1 mM EDTA, 5 mM DTT, and 5% (v/v) glycerol, and consequently eluted three times with a buffer of the same composition but with 1 M NaCl.

Protein was precipitated overnight with 35% (w/v) ammonium sulfate and resuspended in 20 mM Tris-HCl, pH 8, 5% (v/v) glycerol, 0.5 M NaCl, and 5 mM β-mercaptoethanol. The sample was then subject to Ni^2+^-affinity chromatography purification. The eluant protein was dialyzed overnight in a buffer 20 mM Tris-HCl pH 8.0, 5% (v/v) glycerol, 0.1 mM EDTA, 0.5 M NaCl, and 1 mM DTT, and subsequently concentrated and stored at −80°C.

### *In vitro* Transcription Assays

Transcription assays were performed as described previously^37^. Briefly, 50 nM of *Cdiff/Eco* WT or mutant RNAP Eσ^A^ in transcription buffer (10 mM Tris HCl, pH 7.9, 170 mM NaCl, 10 mM MgCl_2_, 1 mM DTT, 5 μg/ml bovine serum albumin (BSA) and 0.1 mM EDTA) was mixed with different concentrations of Fdx (0.01-500 μM). The mixtures were incubated at 37 °C for 5 min to allow for the antibiotic to bind. The *Cdiff rrnC* (GenBank: CP010905.2) dsDNA was added (10nM) to each tube and the samples were incubated for an additional 15 min at 37 °C to allow the formation of the RNAP open complex. Then the dinucleotide (GpU, 20 μM) was added and incubated for 10 min. Transcription was initiated by adding a nucleotide mixture consisting of 20 μM GTP and 1.25 μCi (15 nM)-α-^32^P-GTP. Each reaction was allowed to proceed for 10 min at 37 °C and reactions were quenched by the addition of 2 X stop buffer (0.5X TBE, pH 8.3, 8 M urea, 30 mM EDTA, 0.05% bromophenol blue, and 0.05% xylene cyanol). Reactions were heated at 95°C for 1 min and loaded onto a polyacrylamide gel [20% Acrylamide/Bis acrylamide (19:1), 6M urea, and 1X TBE, pH 8.3]. Transcription products were visualized by using Typhoon FLA 9000 (GE Healthcare) and quantified using ImageQuant software (GE Healthcare). Quantified values were plotted in PRISM and the IC50 was calculated from three independent data sets. The full sequences of the fragments (initiation sites in lowercase, –35 and –10 elements in bold) used for transcription are as follows:

### rrnC

AATAGCTTGTATTAAAGCAGTTAAAATGCATTAATATAGGCTATTTTTATTTTGACA AAAAAATATTTAAAATAAAAGTTAAAAAGTTG**TTGACT**TAGAATAATATAGATGA**T ATTAT**ATATGA**gtg**CCCAAAAGGAGCACCAAAATAAGACAAAAGAACTTTGAAAATT AAACAGTA

### Preparation of WT *Cdiff* Eσ^A^ for cryo-EM

The RNAP core was incubated with 15 molar excess of σ^A^ for 15 min at 37°C and 45 min at 4°C. Then, the complex was purified over a Superose 6 Increase 10/300 GL column (GE Healthcare, Pittsburgh, PA) in gel filtration buffer (20 mM Tris-HCl pH 8.0, 150 mM K-glutamate, 5 mM MgCl_2_, 2.5 mM DTT). The eluted RNAP Eσ^A^ was concentrated to 6 mg/mL (14 μM) by centrifugal filtration (Amicon Ultra). Eσ^A^ was mixed with 100 μM final concentration of Fdx (10 mM stock solution in DMSO) and incubated for 15 minutes at 4 ºC.

### Cryo-EM grid preparation

Before freezing cryo-EM grids, octyl β-D-glucopyranoside was added to the samples to a final concentration of 0.1% ^8^. C-flat holey carbon grids (CF-1.2/1.3-4Au, Protochips, Morrisville, NC) were glow-discharged for 20 sec before the application of 3.5 μL of the samples. Using a Vitrobot Mark IV (Thermo Fisher Scientific Electron Microscopy, Hillsboro, OR), grids were blotted and plunge-froze into liquid ethane with 100% chamber humidity at 22 ºC.

### Cryo-EM data acquisition and processing

*Cdiff* Eσ^A^ with Fdx grids were imaged using a 300 keV Titan Krios (Thermo Fisher Scientific Electron Microscopy) equipped with a K3 Summit direct electron detector (Gatan, Pleasanton, CA). Dose-fractionated movies were recorded in counting mode using Leginon at a nominal pixel size of 1.083 Å/px (micrograph dimensions of 5,760 × 4,092 px) over a nominal defocus range of −1 μm to −2.5 μm^38^. Movies were recorded in “counting mode” (native K3 camera binning 2) with a dose rate of 30 electrons/physical pixel/s over a total exposure of 2 s (50 subframes of 0.04 s) to give a total dose of ∼51 e-/Å^2. A total of 6,930 movies were collected. Dose-fractionated movies were gain-normalized, drift-corrected, summed, and dose-weighted using MotionCor2^39^. The contrast transfer function was estimated for each summed image using Patch CTF module in cryoSPARC v2.15.0^40^. cryoSPARC Blob Picker was used to pick particles (no template was supplied). A total of 2,502,242 particles were picked and extracted from the dose-weighted images in cryoSPARC using a box size of 256 pixels. Particles were sorted using cryoSPARC 2D classification (number of classes, N=50), resulting in 2,415,902 curated particles. Initial models (Ref 1: RNAP, Ref 2: decoy 1, Ref 3: decoy 2) were generated using cryoSPARC *Ab initio* Reconstruction^40^ on a subset of 81,734 particles. Particles were further curated using Ref 1-3 as 3D templates for cryoSPARC Heterogeneous Refinement (N=6), resulting in the following: class1 (Ref 1), 464,460 particles; class2 (Ref 1), 641,091 particles; class3 (Ref 2), 296,508 particles; class4 (Ref 2), 296,203 particles; class5 (Ref 3), 390,575 particles; class6 (Ref 3), 327,065 particles. Particles from class1 and class2 were combined and further curated with another round of Heterogeneous Refinement (N=6), resulting in the following: class1 (Ref 1), 262,185 particles; class2 (Ref 1), 394,040 particles; class3 (Ref 2), 110,023 particles; class4 (Ref 2), 110,743 particles; class5 (Ref 3), 104,013 particles; class6 (Ref 3), 124,547 particles. Curated particles from class 2 were refined using cryoSPARC Non-uniform Refinement ^40^ and then further processed using RELION 3.1-beta Bayesian Polishing^41^. Per-particle CTFs were estimated for the polished particles using cryoSPARC Homogeneous Refinement with global and local CTF refinement enable^40^. These particles were further curated using cryoSPARC Heterogeneous Refinement (N=3), resulting in the following: class1 (Ref 1), 85,470 particles; class2 (Ref 1), 231,310 particles; class3 (Ref 1), 77,250 particles. Particles from class2 were selected for a subsequent cryoSPARC Heterogeneous Refinement (N=3), resulting in the following: class1 (Ref 1), 19,282 particles; class2 (Ref 1), 182,390 particles; class3 (Ref 1), 29,638 particles. Particles in class2 were refined using cryoSPARC Non-uniform Refinement^42^ resulting in final 3D reconstruction containing 182,390 particles with nominal resolution of 3.26 Å. Local resolution calculations were generated using blocres and blocfilt from the Bsoft package^43^.

### Model building and refinement

A homology model for *Cdiff* Eσ^A^ was derived using SWISS-MODEL^44^ and PDBs: 5VI5^20^ for αI and αII; 6BZO^7^ for β, β’, and σ^A^; and 6FLQ^45^ for ω. The homology model was manually fit into the cryo-EM density maps using Chimera^46^ and rigid-body and real-space refined using Phenix^47^. A model of Fdx was used from the previous structure PDB ID: 6BZO to place in the cryo-EM map^7^. Rigid body refinement for rigid domains of RNAP was performed in PHENIX. The model was then manually adjusted in Coot^48^ and followed by all-atom and B-factor refinement with Ramachandran and secondary structure restraints in PHENIX. The BBM2 modules were manually built using the *Eco*^16^ (PDB ID:4LK1) and *Bsub* structures ^17^ (PDB ID:6ZCA) as a template. The refined model was ‘shaken’ by introducing random shifts to the atomic coordinates with RMSD of 0.163 Å in phenix.pdbtools^47^. The shaken model was refined into half-map1 and FSCs were calculated between the refined shaken model and half-map1 (FSChalf1 or work), half-map2 (FSChalf2 or free, not used for refinement), and combined (full) maps using phenix.mtriage^49^. Unmasked log files were plotted in PRISM and the FSC-0.5 was calculated for the full map.

### Reporting summary

Further information on research design is available in the Nature Research Reporting Summary linked to this paper.

### Data and materials availability

Cryo-EM maps and atomic models have been deposited in the Electron Microscopy Database (EMDB accession codes 23210) and the Protein Database (PDB accession codes 7L7B). Unique materials are available from the corresponding authors on request.

## Acknowledgments

We thank Seth Darst for helpful discussions during this study. We thank Rachel Mooney for providing σ ^70^ protein for transcription assays. We thank M. Ebrahim, J. Sotiris, and Honkit Ng at The Rockefeller University Evelyn Gruss Lipper Cryo-electron Microscopy Resource Center. Some of this work was performed at the Simons Electron Microscopy Center and National Resource for Automated Molecular Microscopy located at the New York Structural Biology Center, supported by grants from the Simons Foundation (SF349247), NYSTAR, the Agouron Institute (F00316), and the NIH (GM103310, OD019994). We thank Ed Eng and Kashyap Maruthi for collecting cryo-EM data. This research was supported by grants from the NIH to R.L. (GM38660) and E.A.C. (GM114450).

## Author contributions

E.A.C. and R.L. supervised this work. X.C. and H.B. carried out biochemical and functional assays. H.B. and J.C. determined the cryo-EM structures. X.C. performed bioinformatic analysis.

Y.B. assisted with protein purifications. X.C., H.B., E.A.C., and R.L. wrote the manuscript with input from all authors.

## Additional information

**Supplementary Information** is available for this paper.

**Correspondence and requests for materials** should be addressed to Elizabeth A. Campbell and Robert Landick.

## Extended data figures and tables

**Extended Data Fig. 1.**
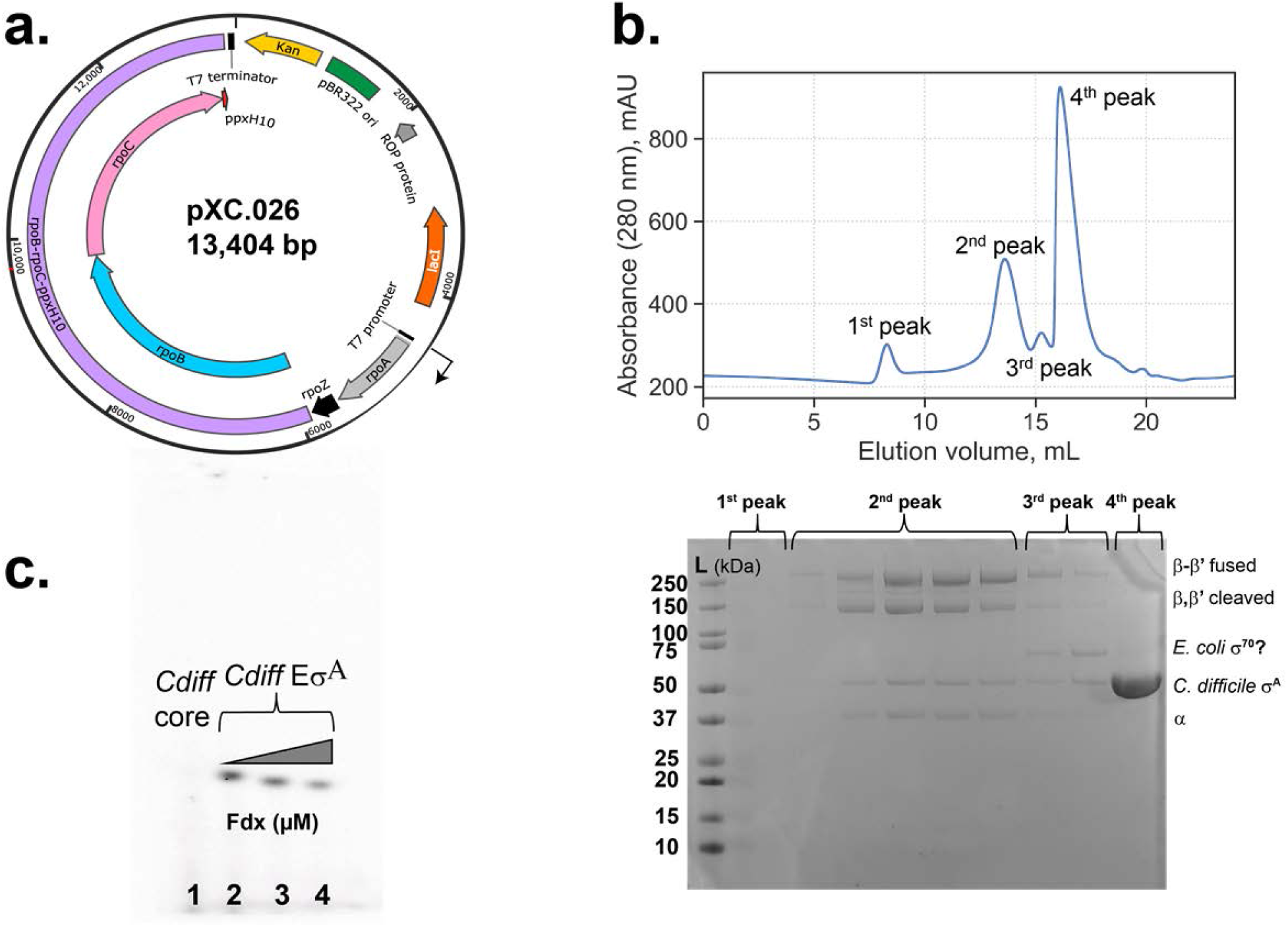
Overexpression and purification of *Cdiff* RNAP. **a**, pXC.026, overexpression plasmid for the *Cdiff rpoA, rpoZ, rpoB*, and *rpoC* genes (encoding the α, ω, β, and β′ subunits of *Cdiff* RNAP, respectively). The β and β′ subunits were fused with an inter-subunit 10-amino-acid (aa) linker (LARHVGGSGA) and a C-terminal Rhinovirus 3C protease-cleavable His_10_ tag. **b**, (Top) Size-exclusion chromatography profile for the assembled *Cdiff* RNAP Eσ^A^. (Bottom) Coomassie-stained SDS-PAGE of individual fractions from major peaks. RNAP subunits are labeled on the right of the gel. *Cdiff* RNAP for biochemistry and structural biology was taken from pooled fractions of the second peak. **c**, Abortive transcription assay with *Cdiff* core and Eσ^A^ using the *Cdiff rrnC* promoter as DNA template. The transcriptional activity of *Cdiff* Eσ^A^ was inhibited with increasing concentrations of Fdx. Lane 1, *Cdiff* RNAP core; lane 2, *Cdiff* Eσ^A^; lane 3, *Cdiff* Eσ^A^ with 0.2 μM Fdx added; lane 4, *Cdiff* Eσ^A^ with 2 μM Fdx added.

**Extended Data Fig. 2.**
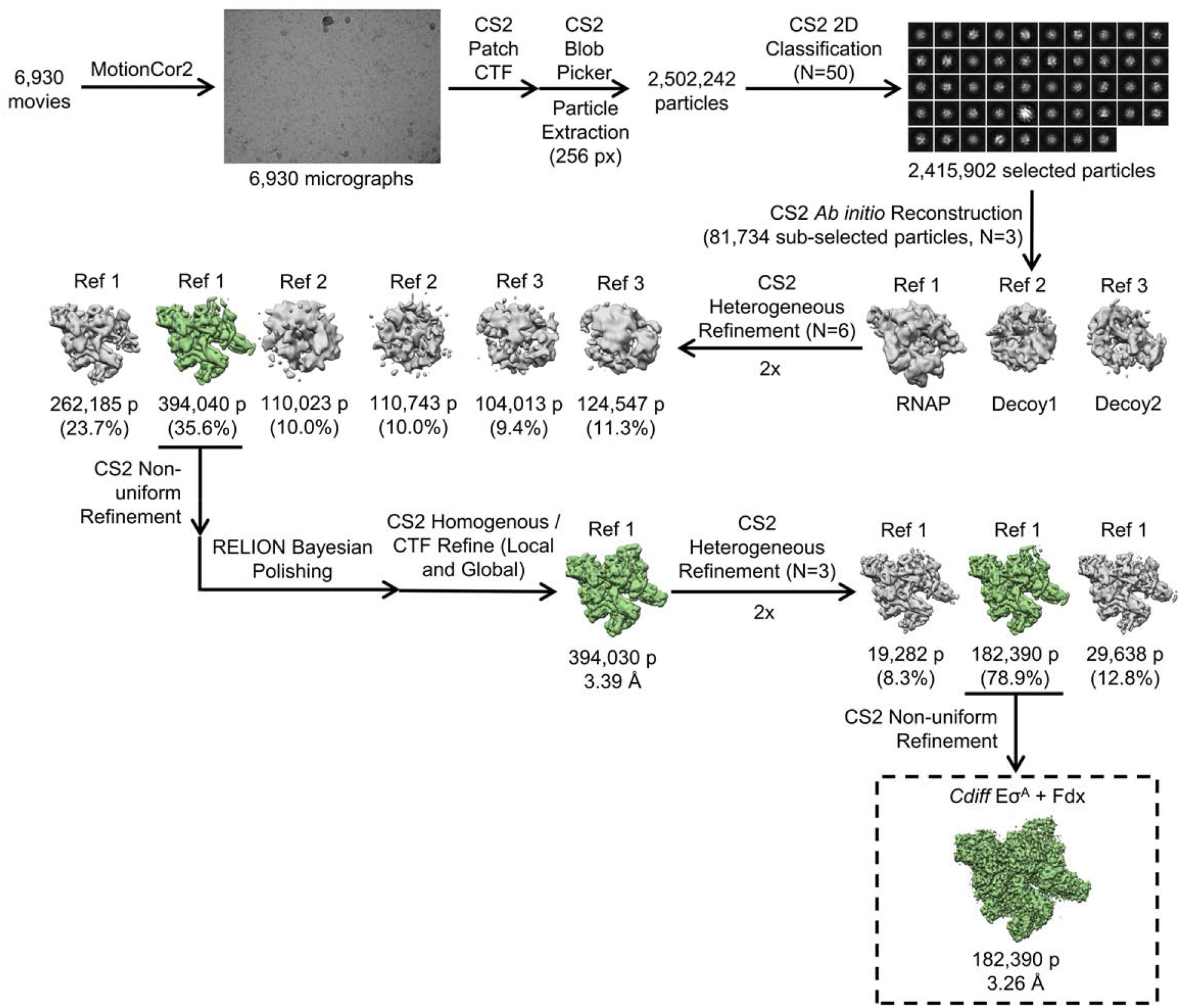
Cryo-EM processing pipeline. Flow chart showing the image-processing pipeline for the cryo-EM data of *Cdiff* Eσ^A^/Fdx complexes, starting with 6,930 dose-fractionated movies collected on a 300-keV Titan Krios (FEI) equipped with a K3 Summit direct electron detector (Gatan). Movies were frame-aligned and summed using MotionCor2^39^. CTF estimation for each micrograph was calculated with cryoSPARC2^40^. A representative micrograph is shown following processing by MotionCor2^39^. Particles were auto-picked from each micrograph with cryoSPARC2 ^40^ Blob Picker and then sorted by 2D classification using cryoSPARC2 to assess quality. The selected classes from the 2D classification are shown. After picking and cleaning by 2D, the dataset contained 2,415,902 particles. A subset of particles was used to generate an *ab initio* templates in cryoSPARC2 and 3D heterogeneous refinement was performed with these templates using cryoSPARC^40^. One major, high-resolution class emerged, which was polished using RELION^41^ and further cleaned with two more 3D heterogenous refinements. The final 182,390 particles were refined using cryoSPARC Non-Uniform refinement^42^.

**Extended Data Fig. 3.**
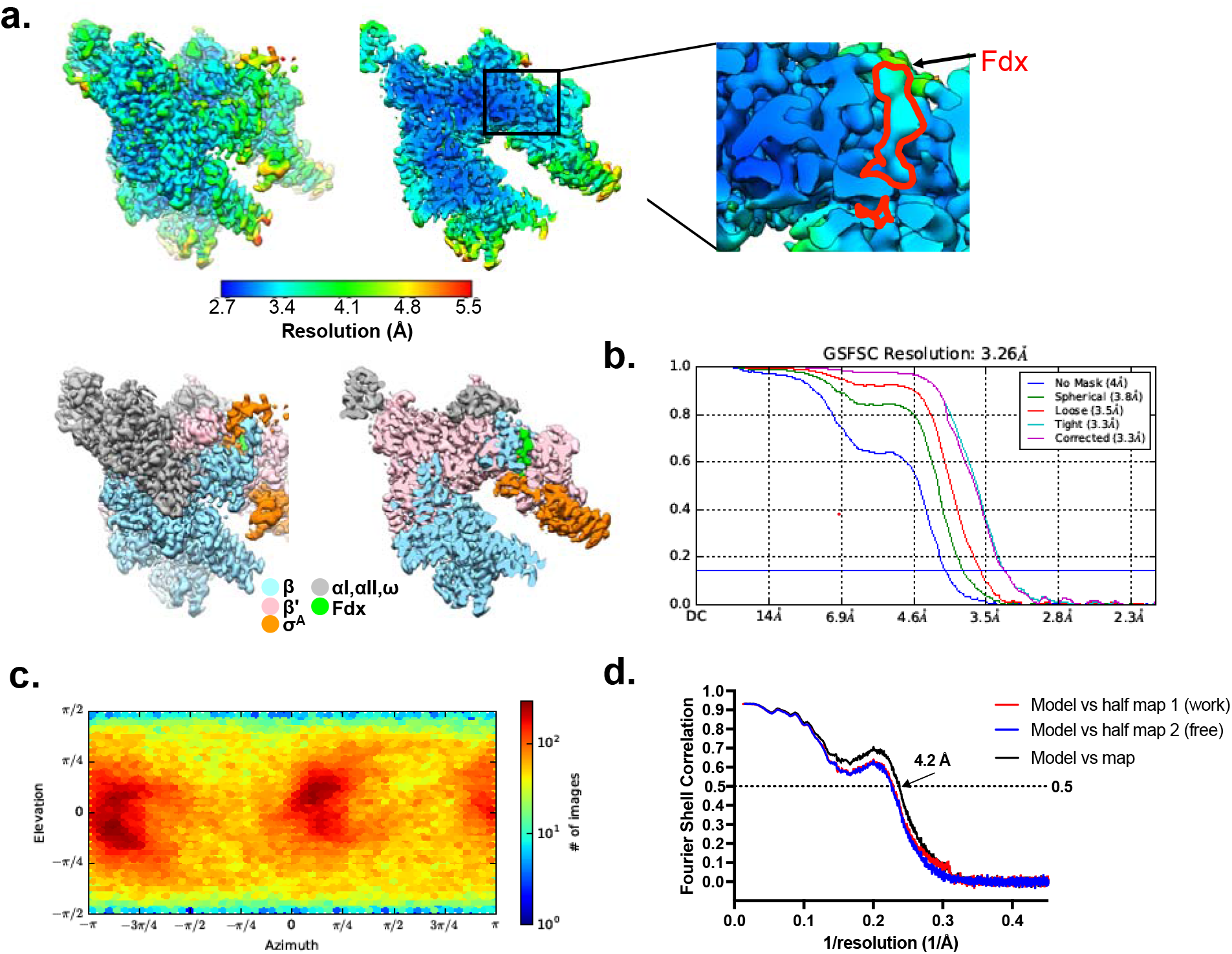
Cryo-EM analysis. **a**, Top left, the 3.26 Å-resolution cryo-EM density map of *Cdiff* Eσ^A^/Fdx. Top right, a cross-section of the structure, showing the Fdx. Bottom, same views as above, but colored by local resolution, The boxed region is magnified and displayed as an inset. Density for Fdx is outlined in red.^15^. **b**, Gold-standard FSC plots of the *Cdiff* Eσ^A^/Fdx complex from cryoSPARC^40^. The dotted line represents the gold-standard 0.143 FSC cutoff which indicates a nominal resolution of 3.26 Å. **c**, Angular distribution calculated in cryoSPARC for *Cdiff* Eσ^A^ /Fdx particle projections. Heat map shows number of particles for each viewing angle (less = blue, more = red)^40^. **d**, Cross-validation FSC plots for map-to-model fitting were calculated between the refined structure of *Cdiff* Eσ^A^/Fdx and the half-map used for refinement (work, red), the other half-map (free, blue), and the full map (black). The dotted black line represents the 0.5 FSC cutoff determined for the full map^49^.

**Extended Data Fig. 4.**
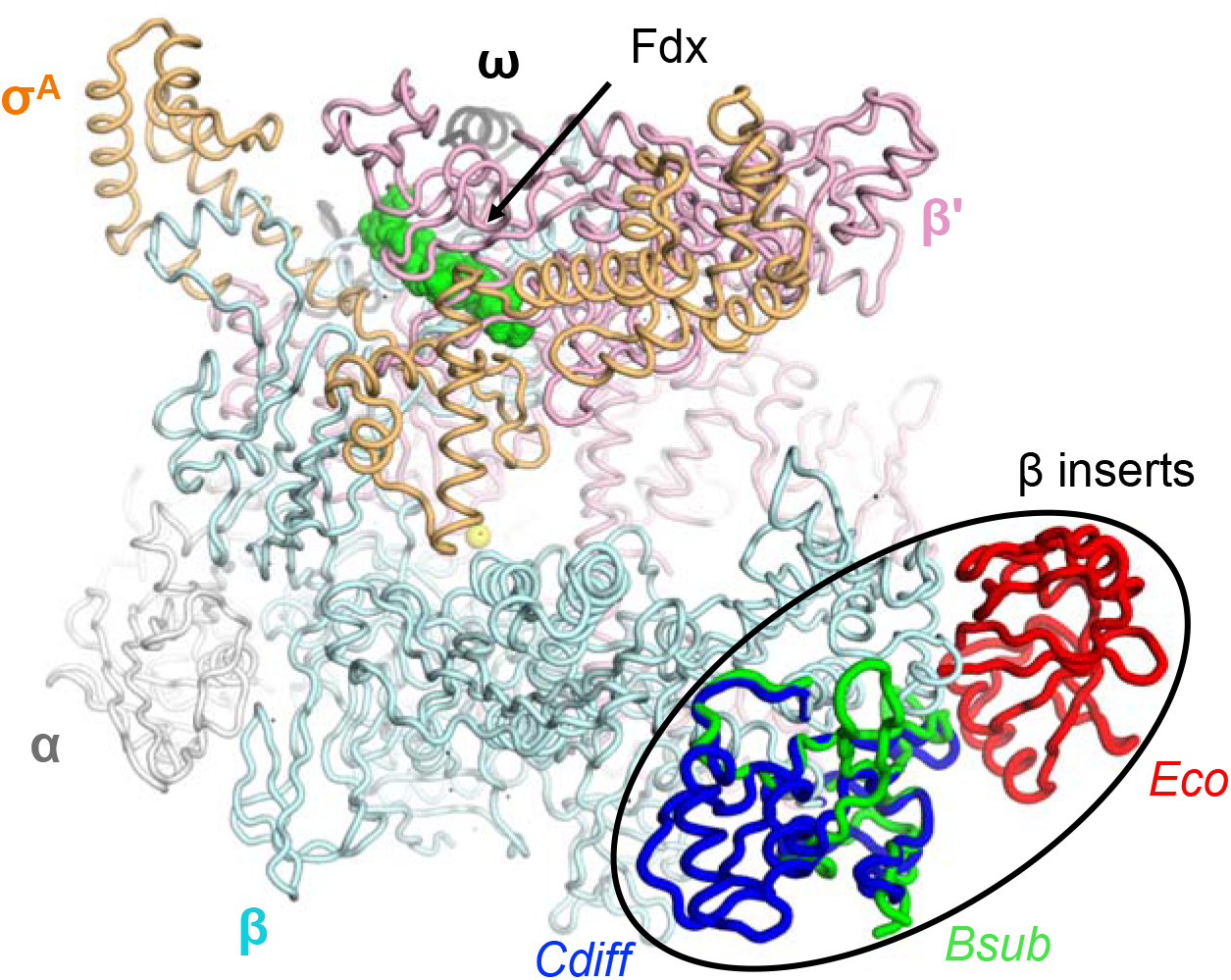
Differences between *Cdiff* and other bacterial RNAPs. The lineage-specific β inserts are shown for *Cdiff* RNAP in dark blue, *Eco* RNAP in red^16^ (PDB ID:4LK1), *Bsub* RNAP in green^17^ (PDB ID:6ZCA). The Fdx is shown in green spheres, and the active site Mg^2+^ is shown as a yellow sphere. Superimposition of the RNAPs from each organism was performed in PyMOL. Only the *Cdiff* Eσ^A^ is shown.

**Extended Data Fig. 5.**
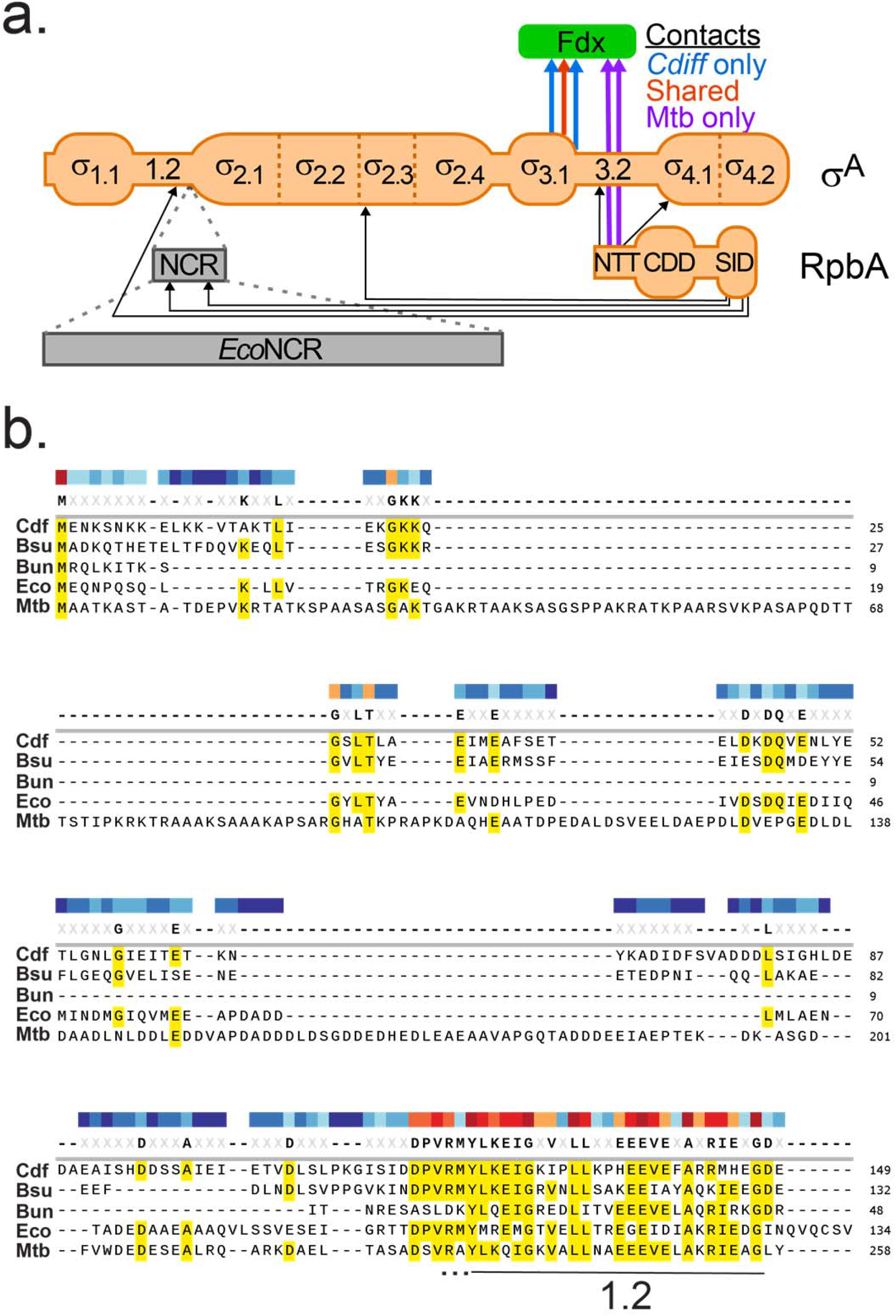
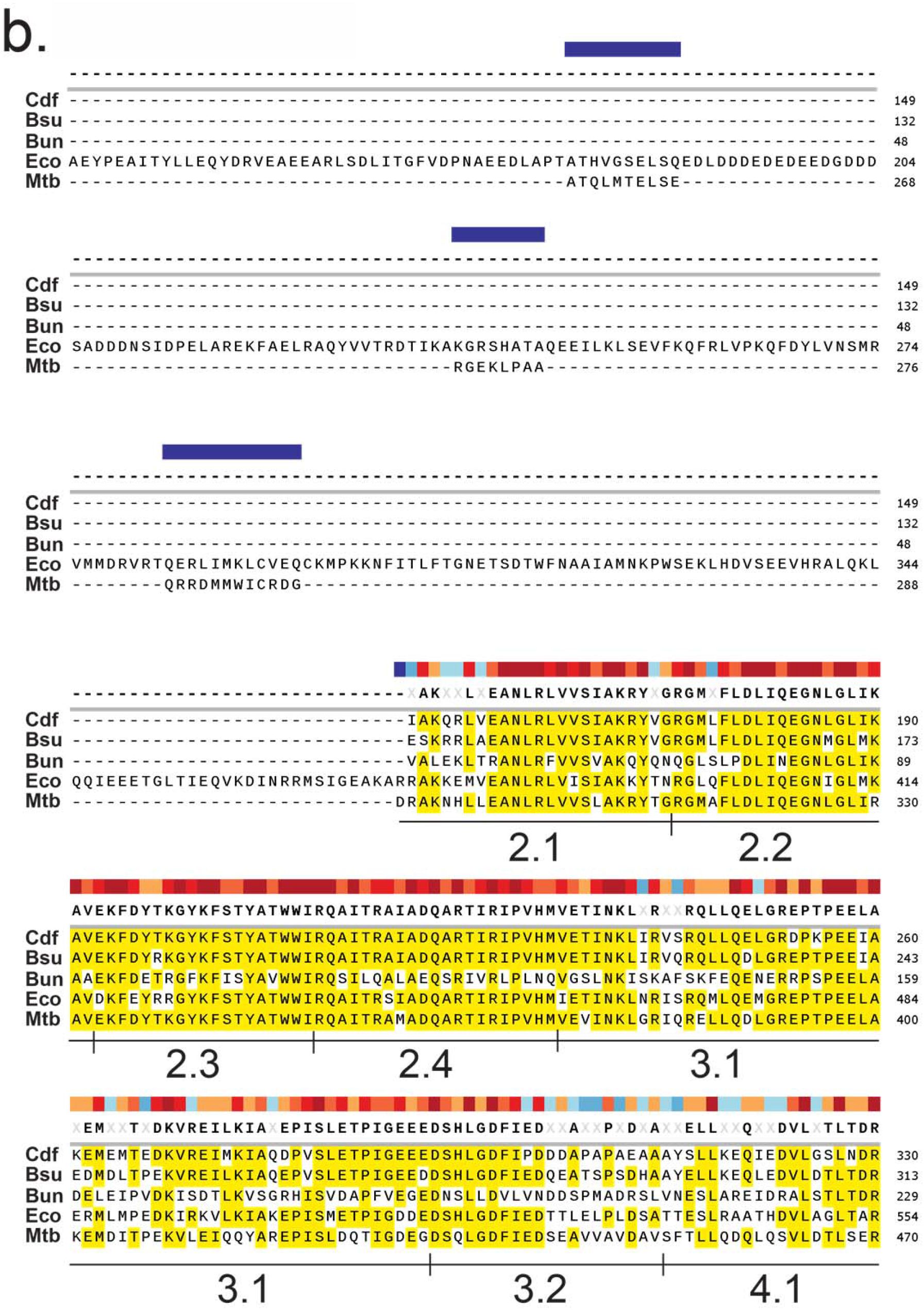
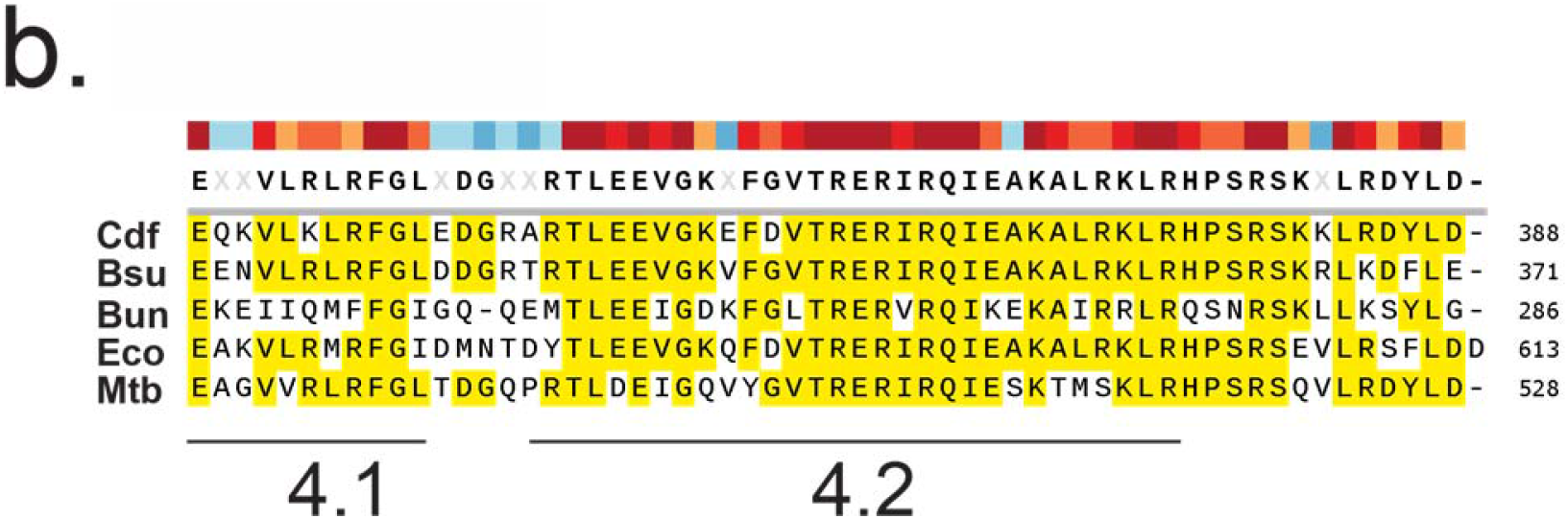
Differences in σ^A^–Fdx contacts between *Cdiff* and *Mtb* and σ^A^ sequence alignment. **a**, Conserved regions of *Cdiff* σ^A^ compared to *Mtb* σ^A^ and *Eco* σ^70^. *Mtb* σ^A^ has a much shorter σ^A^ NCR than *Eco* σ^70^, but the residues in the short *Mtb* NCR that contact RbpA are not present in either *Cdiff* or *Eco*^50^. *Mtb* RbpA contacts Fdx whereas *Cdiff* σ^A^ makes more contacts to Fdx than does *Mtb* σ^A^. Black arrows indicate RpbA-σ^A^ contacts whereas colored arrows indicate Fdx contacts to σ^A^ and RpbA, which includes one shared contact between *Mtb* and *Cdiff* σ^A^ (red arrow). **b**, Amino acid-sequence alignment of σ^A^ for diverse representatives of bacteria species. Identical residues are highlighted in yellow. Gaps are indicated by dashed lines. Conserved σ regions are labeled underneath the alignment. The three letter species code is as follows: Cdf, *Clostridioides difficile*; Bsu, *Bacillus subtilis*; Bun, *Bacteroides uniformis*; Eco, *Escherichia coli*; Mtb, *Mycobacterium tuberculosis*.

**Extended Data Fig. 6.**
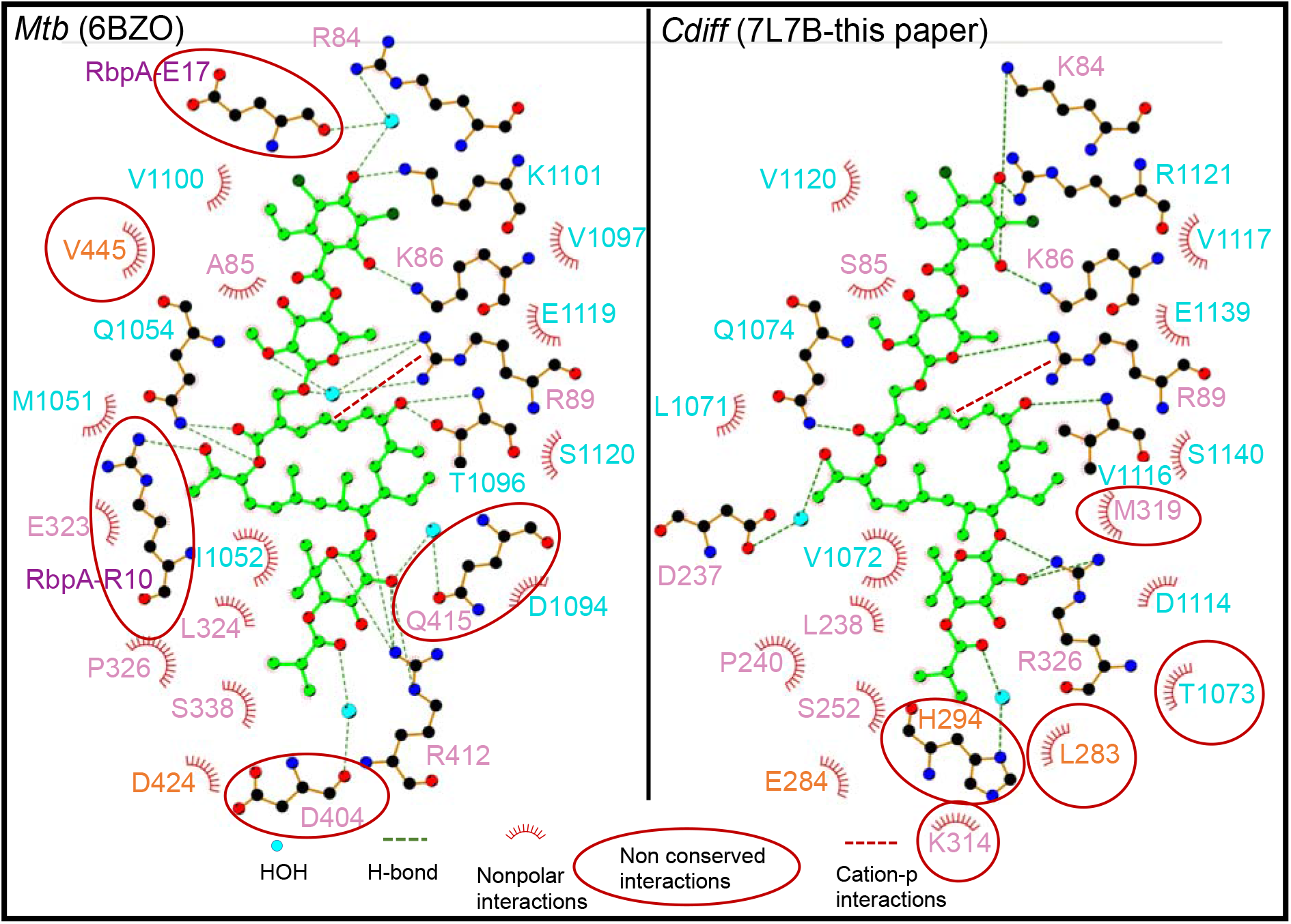
Fdx binding residues in *Mtb* RbpA-Eσ^A^ and *Cdiff* Eσ^A^. Ligplot^51^ was used to determine contacts between Fdx and *Mtb* RbpA-Eσ^A^ (left) and *Cdiff* E**σ**^**A**^ (right). Cyan sphere, H_2_O; green dashed line, hydrogen bond or salt bridge; red arc, van der Waals interactions; red dashed line, cation-π interactions. Note that in ligplot of the *Cdiff* Eσ^A^/Fdx interactions, V1143 (discussed in the text as one of the residues when mutated cause Fdx-resistance) did not make the distance cutoff (4.5 Å) as it was located 4.7 Å away from Fdx. The RNAP β, β’ and σ^A^ residues are in cyan, pink, and orange respectively. The two *Mtb* RbpA residues (E17, R10) that interact with Fdx are colored in purple and indicated in the text. The Fdx-interacting residues that do not have corresponding interactions between *Cdiff* and *Mtb* are highlighted in red circles.

**Extended Data Fig. 7.**
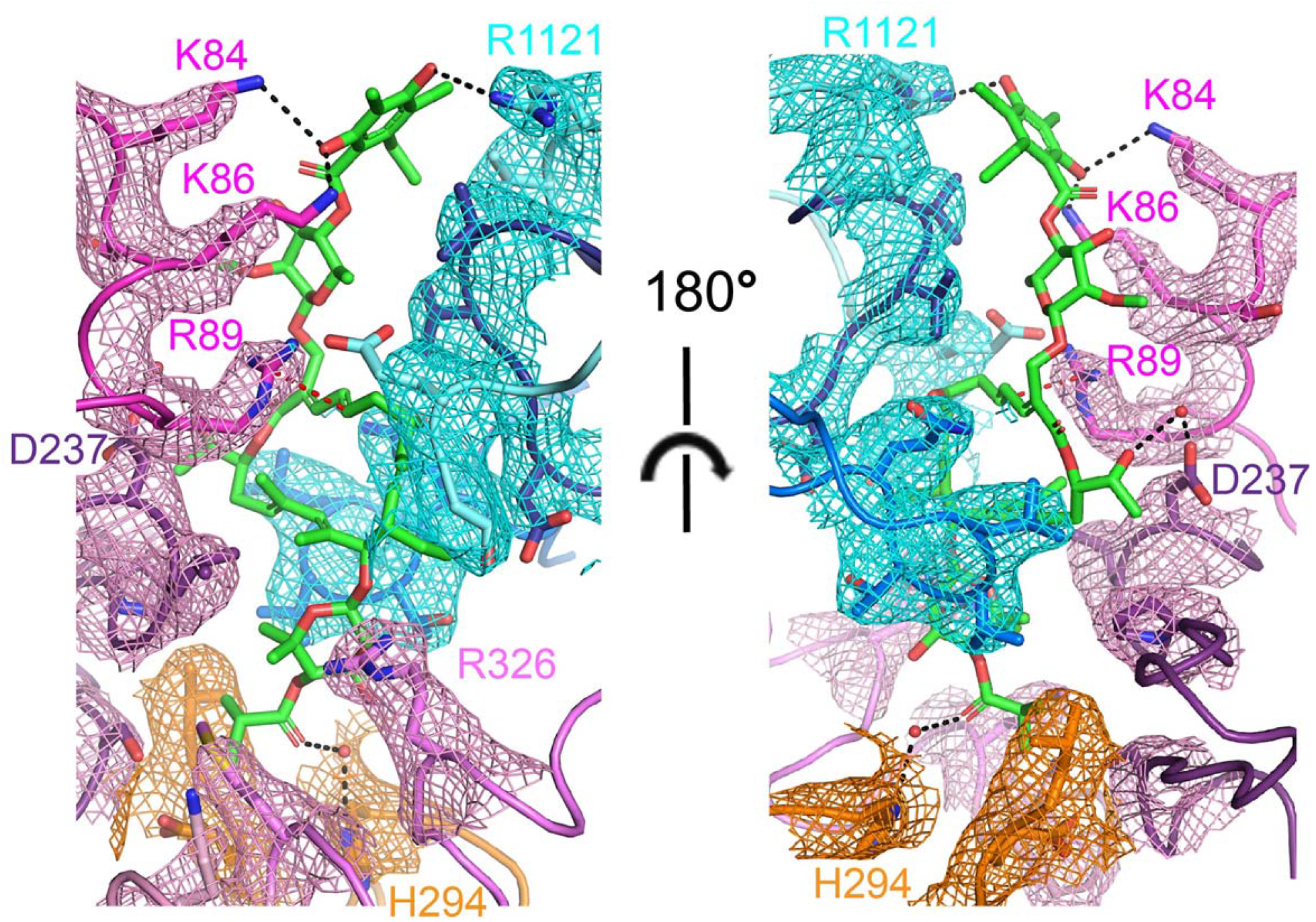
The cryo-EM density map of residues interacting with Fdx. Coloring of the residues is consistent with RNAP subunits coloring in Fig. 3, the stick model and cryo-EM densities are color-coded as follows: Pink: β-subunit, cyan: β’-subunit, and orange: σ^A^. Water molecules are shown as red spheres. The residues that form hydrogen bonds (black dotted line) with Fdx are labeled.

**Extended Data Fig. 8.**
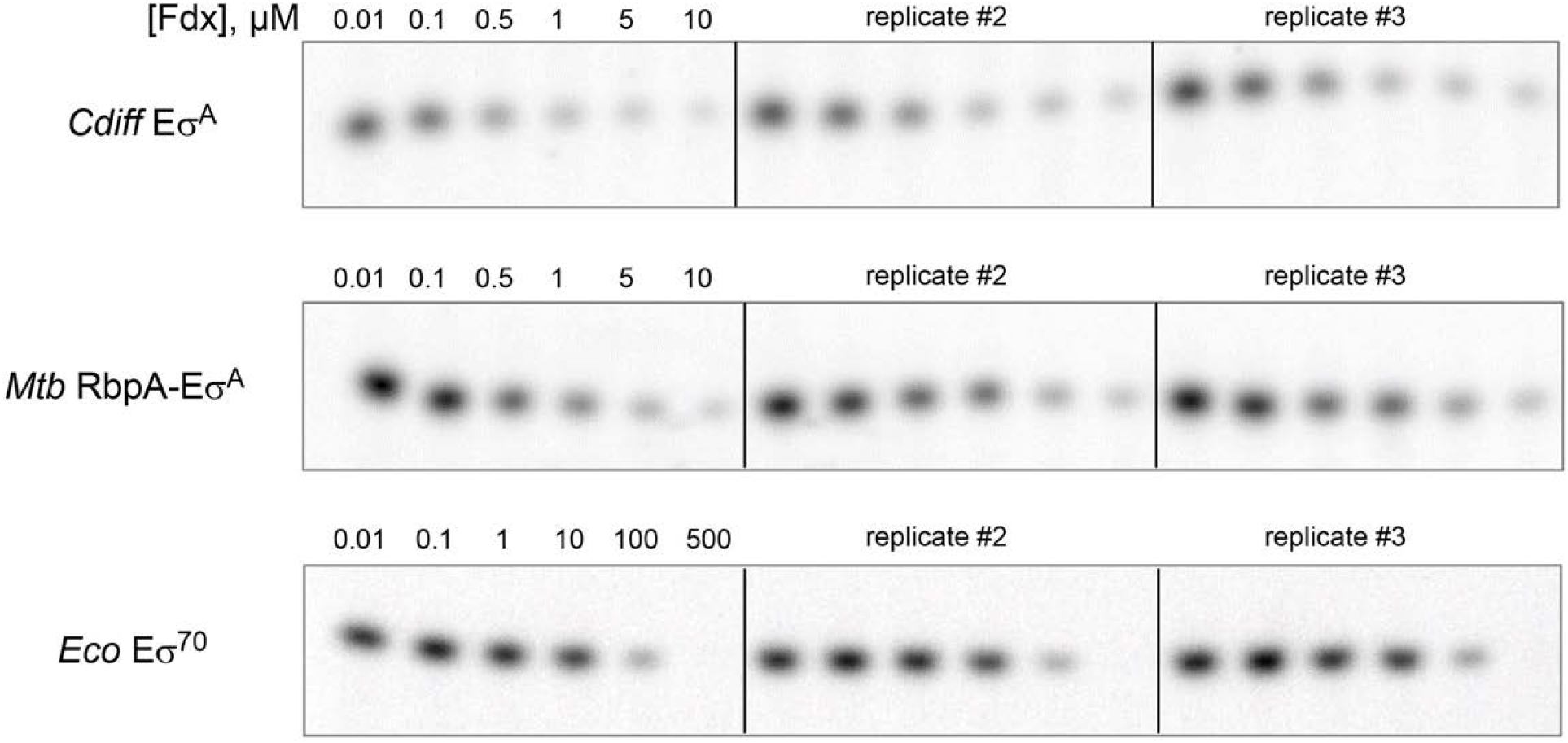
*In vitro* abortive transcription assays used to determine Fdx IC50 of *Cdiff* and *Mtb* Eσ^A^and *Eco*Eσ^70^. Data is plotted in Fig. 2D. Abortive ^32^P-RNA products (G_p_U_p_G) synthesized on *Cdiff* rrnC promoter were quantified in the presence of increasing concentrations of Fdx. For each Eσ^A^ (or Eσ^70^), three independent experiments were performed and analyzed on the same gel.

**Extended Data Fig. 9.**
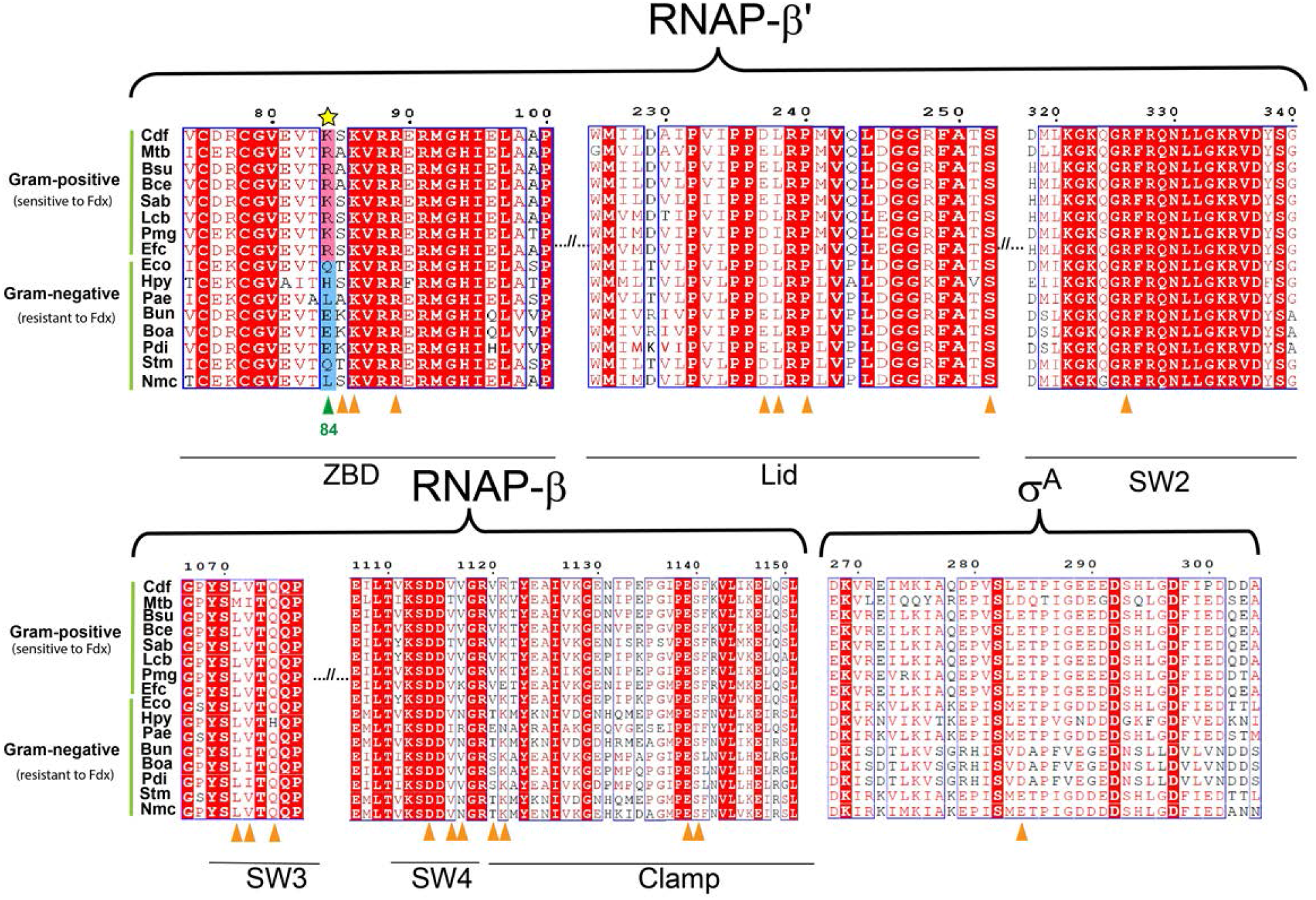
Comparative sequence alignment of key structural components of RNAPs that interact with Fdx between Fdx-resistant and sensitive bacteria. The Fdx interacting regions are labeled on the top of sequence alignment. Locations of residues contacting Fdx in both *Cdiff* and *Mtb* are labeled by triangles underneath sequences. For gram-positive bacteria that are sensitive to Fdx, the corresponding residue at *Cdiff* β’K84 is either K or R, which is highlighted in pink background. For gram-negative bacteria that are resistant to Fdx, the residue at β’K84 is neutral Q, L, or negative E, which is highlighted in blue background. Conserved residues are shown as white letters on a red background, and similar residues are shown as red letters in blue boxes. Cdf, *Clostridioides difficile*; Mtb, *Mycobacterium tuberculosis*; Bsu, *Bacillus subtilis*; Bce, *Bacillus cereus*, Sab, *Staphylococcus aureus*; Lcb, *Lactobacillus casei*; Pmg, *Peptococcus magna;* Efc, *Enterococcus faecium*; Eco, *Escherichia coli*; Hpy, *Helicobacter pylori*; Pae, *Pseudomonas aeruginosa*; Bun, *Bacteroides uniformis*; Boa, *Bacteroides ovatus;* Pdi, *Parabacteroides distasonis*; Stm, *Salmonella Choleraesuis*; Nmc, *Neisseria meningitidis*.

**Extended Data Fig. 10.**
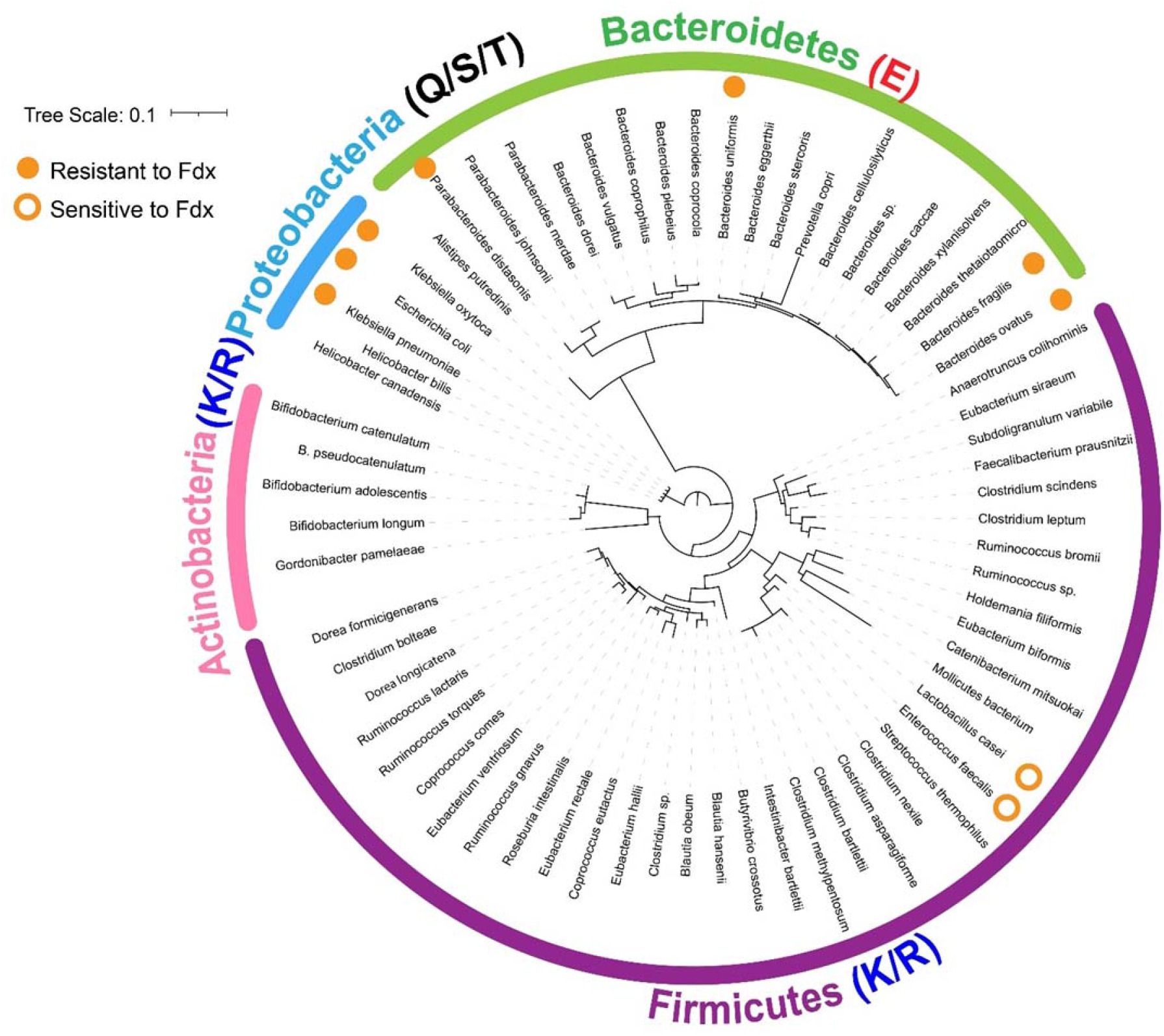
Phylogenetic tree demonstrating the clade-specific distribution of the identity of the Fdx-sensitizer. The tree displays the identity of the amino acid corresponding to position β’K84 of *Cdiff* in the most common species from human gut microbiota. Bacterial species were largely picked from ^28^ and ^29^. The tree was built from 66 small subunit ribosomal RNA sequences by using RaxML^52^ and iTol^53^. Species with experimentally confirmed resistance (MIC > 32 εg/mL) and sensitivity (MIC < 0.125 εg/mL) to Fdx are marked with solid and open orange circle respectively^25^. The amino acid sequence at β’K84 position for corresponding bacteria phyla is denoted by capital letters. The detailed bacterial species are listed in table S4.

**Extended Data Fig. 11.**
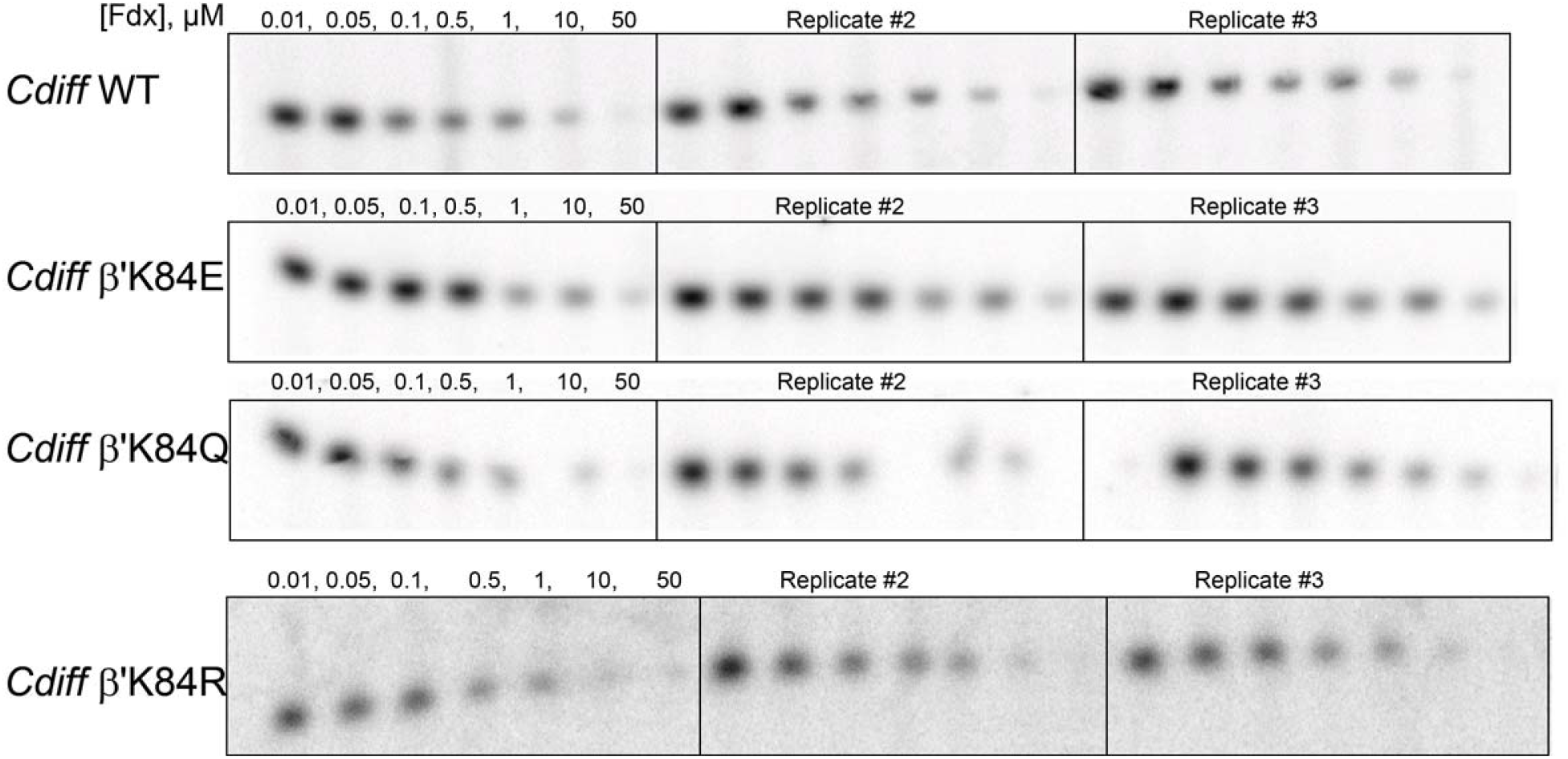
*In vitro* abortive transcription assay measuring Fdx IC50 on *Cdiff* WT and mutant Eσ^A^s related to Fig. 4C. The *Cdiff rrnC* promoter (Fig. 2C) was used as a template. For each RNAP, three independent experiments were performed for and analyzed on the same gel.

**Extended Data Fig. 12.**
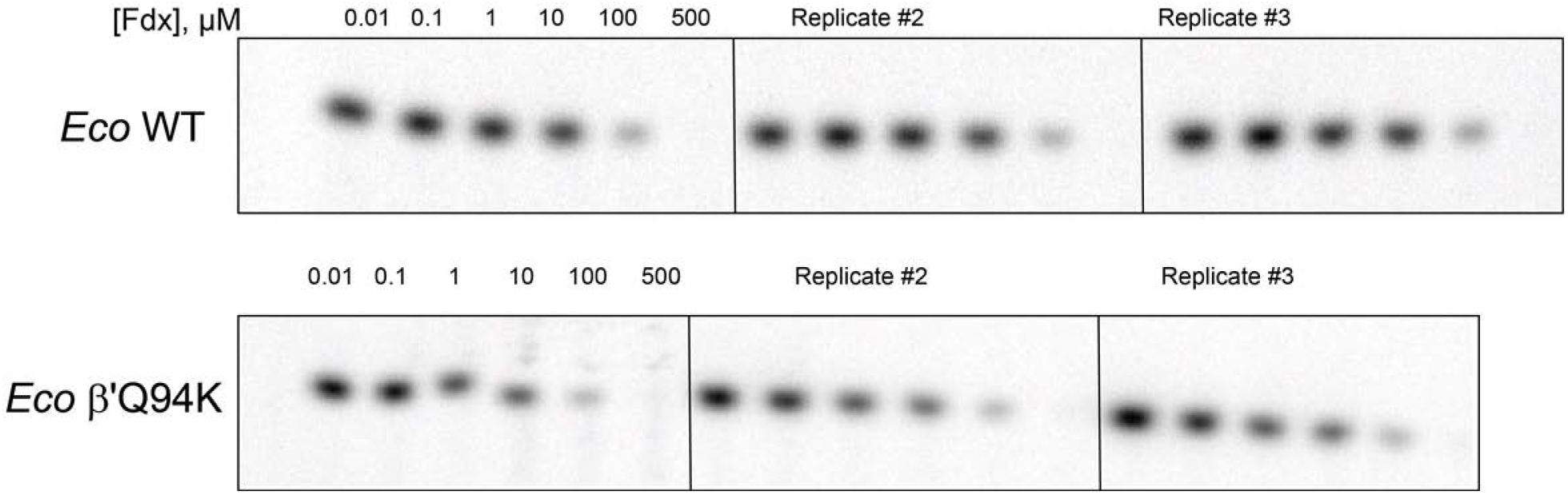
*In vitro* abortive transcription assay measuring Fdx IC50 on *Eco* WT and β’Q94K Eσ^70^s. related to Fig. 4D. The *Cdiff rrnC* promoter (Fig. 2C) was used as a template. For each RNAP, three independent experiments were performed for and analyzed on the same gel.

**Extended Data Table 1.** Antimicrobial efficiency of Fdx on various species.

| Species | MIC (μg/mL) | IC50 (μM)<br>*in vitro* |
| --- | --- | --- |
| *C. difficile* ATCC17857 | 0.031<sup>25</sup> | 0.2 (this study) |
| *M. tuberculosis* (H37Rv) | 0.015<sup>26</sup> | 0.21<sup>7</sup> |
| *E. coli* DH5α | >64<sup>25</sup> | 53<sup>7</sup> |
| *E. coli* D21(*rfa*, *tolC*) | 2<sup>27</sup> | N/A |
| *B. uniformis* ATCC 8492 | >32<sup>25</sup> | N/A |
| *B. ovatus* ATCC8483 | >32<sup>25</sup> | N/A |

**Extended Data Table 2.**
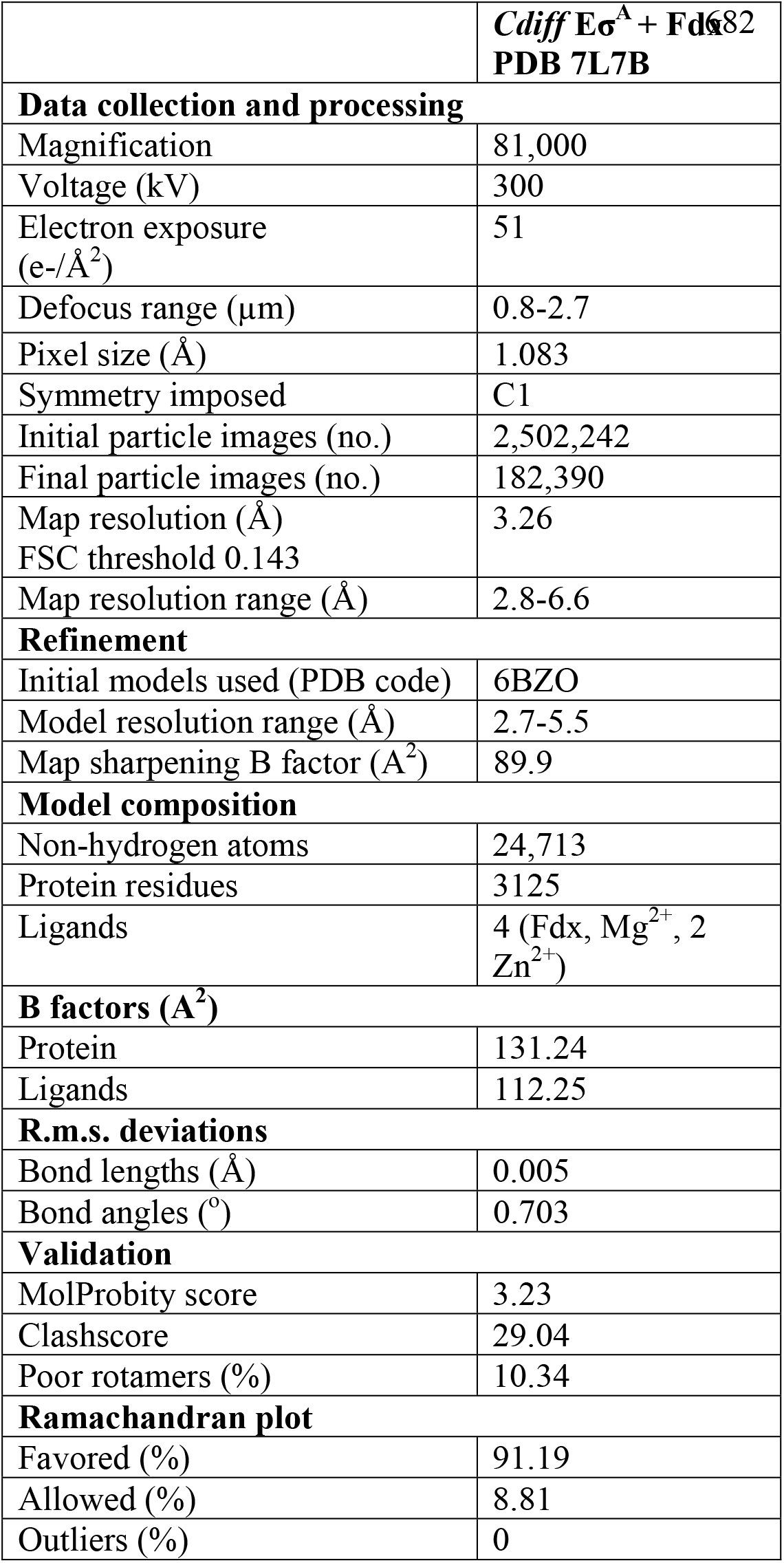
Cryo-EM data collection, refinement, and validation statistics.

**Extended Data Table 3.** Complete list of Fdx interacting residues in *Cdiff* and *Mtb* RNAP Eσ^A^ and comparison for those in *Eco, B. ovatus*, and *B. uniformis*.

| Region | *M. tuberculosis* | *C. difficile* | *E. coli* | *B. ovatus* | *B. uniformis* |
| --- | --- | --- | --- | --- | --- |
| SW3 | β-M1051 ✓ | β-L1071 ✓ | L | L | L |
| SW3 | β-I1052 ✓ | β-V1072 ✓ | I | I | I |
| SW3 | β-Q1054 ✓ | β-Q1074 ✓ | * | * | * |
| SW4 | β-D1094 ✓ | β-D1114 ✓ | * | * | * |
| SW4 | β-T1096 ✓ | β-V1116 ✓ | V | V | V |
| SW4 | β-V1097 ✓ | β-V1117 ✓ | N | V | V |
| Clamp | β-V1100 ✓ | β-V1120 ✓ | T | S | S |
| Clamp | β-K1101 ✓ | β-R1121 ✓ | K | K | K |
| Clamp | β-E1119 ✓ | β-E1139 ✓ | * | * | * |
| Clamp | β-S1120 ✓ | β-S1140 ✓ | * | * | * |
| ZBD | $\beta'$-R84=HOH ✓ | $\beta'$-K84 ✓ | Q | E | E |
| ZBD | $\beta'$-A85 ✓ | $\beta'$-S85 ✓ | T | K | K |
| ZBD | $\beta'$-K86 ✓ | $\beta'$-K86 ✓ | * | * | * |
| ZBD | $\beta'$-R89 ✓ | $\beta'$-R89 ✓ | * | * | * |
| Lid | $\beta'$-E323 ✓ | $\beta'$-D237=HOH ✓ | D | E | E |
| Lid | $\beta'$-L324 ✓ | $\beta'$-L238 ✓ | * | * | * |
| Lid | $\beta'$-P326 ✓ | $\beta'$-P240 ✓ | * | * | * |
| Lid | $\beta'$-S338 ✓ | $\beta'$-S252 ✓ | * | * | * |
| SW2 | $\beta'$-R412 ✓ | $\beta'$-R326 ✓ | * | * | * |
| σ-linker | σ-D424 ✓ | σ-E284 ✓ | E | D | D |
| RbpA | R-R10 ✓<br>R-E17=HOH ✓ | NA<br>NA | NA<br>NA | NA<br>NA | NA<br>NA |
| SW3 | β-T1053 X | β-T1073 ✓ | * | * | * |
| Clamp | $\beta'$-K400(rotamer) X | $\beta'$-K314 ✓ | * | * | * |
| Clamp | $\beta'$-D404=HOH ✓ | $\beta'$-D318 X | * | * | * |
| SW2 | $\beta'$-L405 X | $\beta'$-M319 ✓ | M | S | S |
| SW2 | $\beta'$-Q415=HOH ✓ | $\beta'$-Q329 X | * | * | * |
| σ-linker | σ-L423 X | σ-L283 ✓ | M | V | V |
| σ-linker | σ-Q434 X | σ-H294=HOH ✓ | H | S | S |
| σ-linker | σ-V445 ✓ | σ-P305 X | E | P | P |

✓- van der wall/hydrophobic ✓- Salt-bridge or H bond ✓- cation-pi X no corresponding interactions; \*, identical residues; grey, similar residues; magenta, distinguished residues

**Extended Data Table 4.** List of 66 common bacterial species in human gut profile, related to Fig. 4A. MICs, when reported, are listed. N/A means Not Applicable due to the absence of published values.

| Species | Phylum | AA at $\beta'$K84 | Published MIC |
| --- | --- | --- | --- |
| *Bacteroides vulgatus* | Bacteroidetes | E | N/A |
| *Bacteroides uniformis* | Bacteroidetes | E | 32 $\mu\text{g/mL}$<sup>25</sup> |
| *Bacteroides sp.* | Bacteroidetes | E | N/A |
| *Bacteroides xylanisolvens* | Bacteroidetes | E | N/A |
| *Bacteroides ovatus* | Bacteroidetes | E | 32 $\mu\text{g/ml}$<sup>25</sup> |
| *Bacteroides thetaiotaomicron* | Bacteroidetes | E | N/A |
| *Bacteroides dorei* | Bacteroidetes | E | N/A |
| *Parabacteroides distasonis* | Bacteroidetes | E | 32 $\mu\text{g/ml}$<sup>25</sup> |
| *Parabacteroides merdae* | Bacteroidetes | E | N/A |
| *Bacteroides caccae* | Bacteroidetes | E | N/A |
| *Bacteroides stercoris* | Bacteroidetes | E | N/A |
| *Bacteroides cellulosilyticus* | Bacteroidetes | E | N/A |
| *Bacteroides eggerthii* | Bacteroidetes | E | N/A |
| *Bacteroides plebeius* | Bacteroidetes | E | N/A |
| *Bacteroides coprocola* | Bacteroidetes | E | N/A |
| *Bacteroides coprophilus* | Bacteroidetes | E | N/A |
| *Bacteroides fragilis* | Bacteroidetes | E | > 32 $\mu\text{g/ml}$<sup>25</sup> |
| *Alistipes putredinis* | Bacteroidetes | E | N/A |
| *Parabacteroides johnsonii* | Bacteroidetes | E | N/A |
| *Prevotella copri* | Bacteroidetes | E | N/A |
| *Anaerotruncus colihominis* | Firmicutes | R | N/A |
| *Blautia hansenii* | Firmicutes | K | N/A |
| *Blautia obeum* | Firmicutes | K | N/A |
| *Butyrivibrio crossotus* | Firmicutes | K | N/A |
| *Catenibacterium mitsuokai* | Firmicutes | R | N/A |
| *Intestinibacter bartlettii* | Firmicutes | K | N/A |
| *Clostridium bolteae* | Firmicutes | K | N/A |
| *Clostridium leptum* | Firmicutes | R | N/A |
| *Clostridium methylpentosum* | Firmicutes | R | N/A |
| *Clostridium scindens* | Firmicutes | K | N/A |
| *Clostridium sp.* | Firmicutes | K | N/A |
| *Clostridium nexile* | Firmicutes | K | N/A |
| *Clostridium asparagiforme* | Firmicutes | K | N/A |
| *Clostridium bartlettii* | Firmicutes | K | N/A |
| *Coprococcus comes* | Firmicutes | K | N/A |
| *Coprococcus eutactus* | Firmicutes | K | N/A |
| *Dorea formicigenerans* | Firmicutes | K | N/A |
| *Dorea longicatena* | Firmicutes | K | N/A |
| *Enterococcus faecalis* | Firmicutes | R | 4 $\mu\text{g/mL}$<sup>25</sup> |
| *Eubacterium(Holdemanella) biforme* | Firmicutes | K | N/A |
| *Eubacterium hallii* | Firmicutes | K | N/A |
| *Eubacterium rectale* | Firmicutes | K | N/A |
| *Eubacterium siraeum* | Firmicutes | R | N/A |
| *Eubacterium ventriosum* | Firmicutes | K | N/A |
| *Faecalibacterium prausnitzii* | Firmicutes | K | N/A |
| *Holdemania filiformis* | Firmicutes | K | N/A |
| *Lactobacillus casei* | Firmicutes | R | 1 µg/mL<sup>25</sup> |
| *Mollicutes bacterium* | Firmicutes | R | N/A |
| *Roseburia intestinalis* | Firmicutes | K | N/A |
| *Ruminococcus bromii* | Firmicutes | R | N/A |
| *Ruminococcus gnavus* | Firmicutes | K | N/A |
| *Ruminococcus lactaris* | Firmicutes | K | N/A |
| *Ruminococcus sp* | Firmicutes | K | N/A |
| *Ruminococcus torques* | Firmicutes | K | N/A |
| *Streptococcus thermophilus* | Firmicutes | R | N/A |
| *Subdoligranulum variabile* | Firmicutes | R | N/A |
| *Bifidobacterium adolescentis* | Actinobacteria | K | N/A |
| *Bifidobacterium catenulatum* | Actinobacteria | K | N/A |
| *Bifidobacterium longum* | Actinobacteria | K | N/A |
| *Bifidobacterium pseudocatenulatum* | Actinobacteria | K | N/A |
| *Gordonibacter pamelaee* | Actinobacteria | R | N/A |
| *Escherichia coli* | Proteobacteria | Q | >32 µg/mL<sup>25</sup> |
| *Klebsiella pneumoniae* | Proteobacteria | Q | >32 µg/mL<sup>25</sup> |
| *Klebsiella oxytoca* | Proteobacteria | Q | 64 µg/mL<sup>25</sup> |
| *Helicobacter bilis* | Proteobacteria | T | N/A |
| *Helicobacter canadensis* | Proteobacteria | S | N/A |

